# Chromosomal instability can favor macrophage-mediated immune response and induce a broad, vaccination-like anti-tumor IgG response

**DOI:** 10.1101/2023.04.02.535275

**Authors:** Brandon H. Hayes, Mai Wang, Hui Zhu, Steven H. Phan, Lawrence J. Dooling, Jason C. Andrechak, Alexander H. Chang, Michael P. Tobin, Nicholas M. Ontko, Tristan Marchena, Dennis E. Discher

## Abstract

Chromosomal instability (CIN), a state in which cells undergo mitotic aberrations that generate chromosome copy number variations, generates aneuploidy and is thought to drive cancer evolution. Although associated with poor prognosis and reduced immune response, CIN generates aneuploidy-induced stresses that could be exploited for immunotherapies. In such contexts, macrophages and the CD47-SIRPα checkpoint are understudied. Here, CIN is induced pharmacologically induced in poorly immunogenic B16F10 mouse melanoma cells, generating persistent micronuclei and diverse aneuploidy while skewing macrophages towards an anti-cancer M1-like phenotype, based on RNA-sequencing profiling, surface marker expression and short-term antitumor studies. These results further translate to *in vivo* efficacy: Mice bearing CIN-afflicted tumors with wild-type CD47 levels survive only slightly longer relative to chromosomally stable controls, but long-term survival is maximized when combining macrophage-stimulating anti-tumor IgG opsonization and some form of disruption of the CD47-SIRPα checkpoint. Survivors make multi-epitope, *de novo* anti-cancer IgG that promote macrophage-mediated phagocytosis of CD47 knockout B16F10 cells and suppress tumoroids *in vitro* and growth of tumors *in vivo*. CIN does not greatly affect the level of the IgG response compared to previous studies but does significantly increase survival. These results highlight an unexpected therapeutic benefit from CIN when paired with maximal macrophage anti-cancer activity: an anti-cancer vaccination-like antibody response that can lead to more durable cures and further potentiate cell-mediated acquired immunity.

## Introduction

Chromosomal instability (CIN) has long been thought to be indicative of poor prognosis and reduced immune cell activity against tumors (Davoli et al., 2017; Vasudevan et al., 2021). CIN is a state of high frequency of chromosome mis-segregation, which often generates micronuclei and can ultimately cause aneuploidy—an abnormal number of chromosomes. CIN-induced genomic heterogeneity can serve as a tumor promotor (Sheltzer et al., 2017) and allow some tumor subpopulations to favor aggression, metastatic potential, immune evasion, and resistance to therapies (Ben-David & Amon, 2019; Chunduri & Storchová, 2019; Vasudevan et al., 2021). However, early-stage CIN also induces anti-cancer vulnerabilities (Cohen-Sharir et al., 2021; Vasudevan et al., 2020), such as proliferation deficits (Wang et al., 2021). These early-stage CIN-afflicted cells have yet to adapt and achieve aneuploidies that favor growth and immune evasion (Tripathi et al., 2019, Vasudevan et al., 2021). CIN in diploid cells that is caused by spindle assembly checkpoint disruption via MPS1 kinase inhibition (MPS1i) further induces a senescence-associated secretory pathway phenotype and upregulation of NF-κB and interferon-mediated pathways, among others, that drive immune clearance of chromosomally aberrant cells (Santaguida et al., 2017; Wang et al., 2021). CIN and ploidy changes also inhibit tumor growth in immunocompetent mice while having little effect in immunocompromised mice (Senovilla et al., 2012; Boilève et al., 2013), which suggests CIN somehow increases immunogenicity.

Recent analyses of The Cancer Genome Atlas (TCGA) showed that highly aneuploid tumors include macrophages that are polarized toward a pro-cancer, M2-like phenotype, among other pro-cancer immune changes (Davoli et al., 2017; Taylor et al., 2018). MPS1i-treated cancer cells have also been reported to escape immune-mediated clearance (Wang et al., 2021), with another study reporting that cancer cells respond to CIN with IL-6-STAT3 signaling (Hong et al., 2022) that protects from CIN-induced cell death, minimizes interferon-related anti-cancer responses, and allows cells to adapt to CIN- and aneuploidy-induced stresses. Two key observations stood out among these studies. First, ploidy changes tend to increase factors that can promote macrophage-mediated phagocytosis (Chao et al., 2010; Krysko et al., 2018). Second, pro-survival signaling amidst CIN leads to an increase of other factors (such as IL-6) that induce a pro-cancer, M2-like phenotype (Fernando et al., 2014). Together, these observations led us to hypothesize that macrophages play a key role in influencing CIN- and aneuploidy-afflicted tumors.

Macrophages have become attractive candidates for immunotherapies due to their ability to phagocytose cancer cells (Alvey et al., 2018; Morrissey et al., 2020). Phagocytosis of ‘self’ cells is generally inhibited by the key macrophage checkpoint interaction between SIRPα on the macrophage and CD47 on the target cell (Oldenborg et al., 2000). CD47 is ubiquitously expressed on all cells, including cancer cells (Willingham et al., 2012). While tumor cell engulfment can be driven to some extent via IgG opsonization by using anti-tumor monoclonal antibodies that bind Fc receptors on macrophages (Alvey et al., 2017; Suter et al., 2021; Tsai & Discher, 2008), this is generally insufficient to eliminate cancers, especially solid tumors. Maximal macrophage-mediated phagocytosis is achieved when CD47-SIRPα signaling is simultaneously disrupted. However, achieving complete tumor rejection is still a major challenge for macrophage-oriented therapies (Ingram et al., 2017; Sockolosky et al., 2016), even after having identified such key factors. Previous studies achieved durable cures in immunocompetent mice, with some acquired immunity benefits, but tumor rejection success is modest and variable (∼20-40%; Dooling et al., 2023; Hayes et al., 2023). Additionally, *in vivo* macrophages often polarize toward tumor-associated macrophage (TAM) phenotypes that tend to correlate with poor clinical prognoses (Cerezo-Wallis et al., 2020; Noy & Pollard, 2014). TAMs have poor phagocytic function and can promote tumor growth and invasion (Georgouli et al., 2019). Cancer cell CIN could create conditions in which macrophages can dominate cancer cell proliferation and even avoid becoming TAM-like.

Inspired by recent advances in exploiting CIN-related anti-cancer vulnerabilities, we hypothesized that early-stage CIN in cancer cells might stimulate higher anti-cancer macrophage activity than more chromosomally stable tumors. Our results provide initial *in vitro* and *in vivo* evidence that CIN indeed favors anti-cancer macrophage activity and consistently leads to durable cures in mice under conditions of maximal phagocytosis. Elimination of these CIN-afflicted solid tumors further drives development of both anti-cancer opsonizing IgGs and enhanced cell-mediated immunity, both of which can help suppress growth against aggressive chromosomally stable tumors.

## Results

### CIN-afflicted tumors skew macrophages towards an anti-cancer phenotype

To investigate the possible effects that cancer cell CIN may have in mediating macrophage immune response, we focused on the poorly immunogenic B16F10 mouse melanoma model. To induce CIN, we treated cells for 24 h with the MPS1 inhibitor reversine (Hong et al., 2022; Kitajiam et al., 2022; Santaguida et al., 2010; Santaguida et al., 2017), and after washout and recovery for 48 h (**Fig. 1A**), we quantified CIN in B16F10 cells, characterized its effects in early-stage tumors, and studied potential effects on macrophages *in vitro* (**Fig. 1A**). MPS1i increased the frequency of micronuclei visible in interphase cells (**Fig. 1B**), as micronuclei are often used as a surrogate for CIN and genome instability (Cohen-Sharier et al., 2020; Crasta et al., 2012; Harding et al., 2017; Mackenzie et al., 2017). Regardless of the MPS1i concentration, we saw >10-fold increases of micronuclei over the cell line’s low basal level (∼1% of cells), and two other MPS1i inhibitors AZ3146 and BAY12-17389 confirm such effects (**Fig. S1A**). Micronuclei-positive cells can persist up to 12 days after treatment (**Fig. S1B**), while control cells maintain the low basal levels. The results suggest pre-treatment with MPS1i can simulate CIN in an experimental context even for 1-2 weeks, which may not typically occur at the same frequency during early tumor growth.

**Figure 1.**
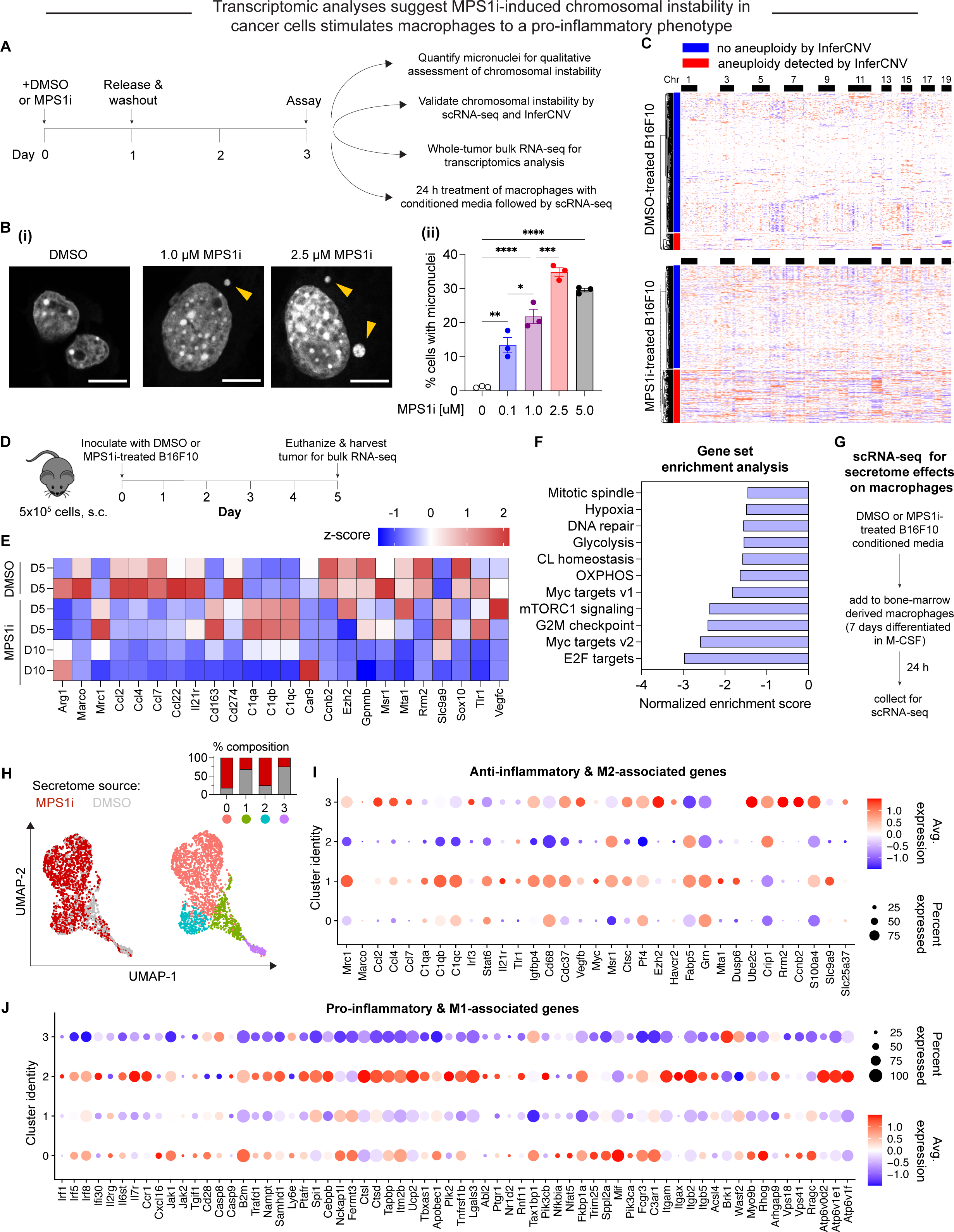
MPS1i-induced chromosomal instability generates microenvironment conditions that favor anti-cancer M1-like macrophages while minimizing skewing to a pro-cancer M2-like phenotype. **(A)** Timeline and schematic for treatment of B16F10 mouse melanoma cells with MPS1 inhibitors (MPS1i). B16F10 cells were treated with MPS1i (reversine) or the equivalent volume of DMSO vehicle control. Cells were treated for 24 h, after which they were washed twice with PBS and allowed to recover for an additional 48 h. After the recovery period elapsed, numerous follow-up experiments were conducted to characterize chromosomal instability, aneuploidy, and effects on BMDMs (BMDMs). **(B) (i)** Representative DNA images of B16F10 cells 72 h after initial treatment, as outlined in (A). Scale bars = 10 μm. Yellow arrowheads point at micronuclei, which are used as a signature readout of chromosomal instability. **(ii)** Quantification of the percentage of B16F10 cells with micronuclei after MPS1i treatment, across a range of different MPS1i concentrations and DMSO vehicle control (mean *±* SEM shown, n = 3 replicates per condition). Statistical significance was calculated by ordinary one-way ANOVA and Tukey’s multiple comparison test (* p < 0.05; ** p < 0.01; *** p < 0.001; **** p < 0.0001). **(C)** Inferred copy number in DMSO and MPS1i-treated B16F10 cells from single-cell RNA-sequencing and the InferCNV pipeline. Cells that are considered aneuploid (labeled as outliers in the InferCNV algorithm) show full-level chromosome gains and/or losses compared to the consensus copy number profile of the DMSO-treated B16F10 population. Approximately 34% of MPS1i-treated B16F10 show aneuploidy, as determined by InferCNV, compared to 10% in the DMSO-treated B16F10 population. Single-cell RNA-sequencing confirms that MPS1i induces chromosomal instability and aneuploidy in B16F10 cells. **(D)** Timeline and schematic for whole-tumor bulk RNA-sequencing. Prior to tumor inoculation, B16F10 CD47 KO cells were treated with 2.5 μM MPS1i (reversine) or the equivalent volume of DMSO vehicle control. Cells were treated for 24 h, after which they were washed twice with PBS and allowed to recover for an additional 48 h. After the recovery period elapsed, all mice were subcutaneously injected with 5x10^5^ B16F10 cells. Five or ten days after tumor inoculation, mice were euthanized, and tumors were excised and disaggregated for bulk RNA-sequencing. **(E)** Heatmap of selected RNA transcripts related to M2 macrophage polarization and pro-tumor function that were differentially expressed in tumors comprised of MPS1i-treated B16F10 compared to tumors comprised of their DMSO-treated counterpart. Heatmap shows log_2_-transformed transcript reads which were then z-score normalized. Generally, M2 and pro-tumor macrophage markers are downregulated at day 5 and even further at day 10 compared to DMSO controls. Tumors analyzed were two comprised of DMSO-treated B16F10 cells five days post-inoculation, two comprised of MPS1i-treated B16F10 cells five days post-inoculation, and two comprised of MPS1i-treated B16F10 cells ten days post-inoculation. **(F)** Top downregulated hallmark gene sets in tumors comprised of MPS1i-treated B16F10 cells, showing downregulated DNA repair, cell cycle, and growth pathways, consistent with observations of slowed growth in culture and in vivo – as subsequently quantified. Gene set enrichment analysis pathways were obtained from MSigDB. Abbreviations: CL homeostasis = cholesterol homeostasis; OXPHOS = oxidative phosphorylation. **(G)** Scheme for conditioned media (secretome) treatment of BMDMs and subsequent characterization by single-cell RNA-sequencing. Media from DMSO and MPS1i-treated B16F10 cells was collected and added 1:1 with fresh media to 7-day differentiated BMDMs (with 20 ng/mL M-CSF). After 24 h, BMDMs were processed for single-cell RNA-sequencing. **(H) (Left)** UMAP plots of expression profiles for all analyzed BMDMs, treated with the secretome from DMSO-treated (gray) or MPS1i-treated (red) B16F10 cells. Each circle represents an individual cell. **(Right)** Same UMAP plots but colors now represent cells clustered together based on similarity of global gene expression. **(Inset)** Composition of each cluster. **(I)** Dot plot showing proportion of cells in each cluster expressing anti-inflammatory and M2-like polarization-associated genes (pro-cancer gene set overall). Indicated to the right are a heatmap scale for average gene expression and a dot size reference for what proportion of cells in the cluster express a gene. Clusters 1 and 3, both of which consist of ∼75% of BMDMs treated with conditioned media from DMSO-treated B16F10, generally show downregulated transcript levels for genes associated with skewing macrophages to a pro-inflammatory, anti-cancer, M1-like phenotype. Cluster 3, in particular, shows significant downregulated. On the contrary, clusters 0 and 2, which consist of 75% of BMDMs treated with conditioned media from MPS1i-treated B16F10, show upregulated expression of many of these anti-cancer genes. Cluster 2, in particular, shows signatures of high upregulation. **(J)** Dot plot showing proportion of cells in each cluster expressing pro-inflammatory and M1-like polarization-associated genes (anti-cancer gene set overall). Indicated to the right are a heatmap scale for average gene expression and a dot size reference for what proportion of cells in the cluster express a gene. Clusters 1 and 3, both of which consist of ∼75% of BMDMs treated with conditioned media from DMSO-treated B16F10, generally show upregulated transcript levels for genes associated with skewing macrophages to an anti-inflammatory, pro-cancer, M2-like phenotype. On the contrary, clusters 0 and 2, which consist of 75% of BMDMs treated with conditioned media from MPS1i-treated B16F10, show either little-to-no expression or downregulation of many of these genes.

Upon confirmation of increased micronuclei induction, we then proceeded to quantitatively confirm and assess copy number variations that resulted from MPS1i treatment using single-cell RNA-sequencing. For DMSO-treated B16F10 cells, approximately 10% of the population were considered aneuploid outliers (**Fig. 1C-top; S1C-D**) – cells with copy number profiles that are 2.5 standard deviations away from the distribution peak (see **Methods**). However, 34% of the MPS1i-treated B16F10 cells were aneuploid (**Fig. 1C-bottom; S1C-D**).

After confirming that MPS1i indeed induces CIN and quantifiable aneuploidy in B16F10 cells, we next sought to understand how CIN may affect early-stage immune response and tumor development. We proceeded to establish tumors in mice with either MPS1i-treated or DMSO-treated B16F10 (following the same schema outlined in **Fig. 1A**). We isolated tumors from mice at two timepoints, five and ten days after initial challenge, and then we processed whole-tumors for bulk RNA-sequencing (**Fig. 1D**). Comparison of tumors comprised of MPS1i-treated and DMSO-treated cells by differential gene expression showed distinct downregulated transcripts for numerous genes encoding M2-like (pro-cancer) macrophage polarization markers (**Fig. 1E**). At day 5 post-challenge, classical M2 markers *Arg1*, *Marco*, and *Cd274* are downregulated, and although several M2 markers are upregulated (*Mrc1*, *Cd163*, and C1q complement genes), by day 10, these, *Marco*, and *Cd27* are all downregulated. Furthermore, at both days 5 and 10, cytokines *Ccl2*, *Ccl4*, *Ccl7*, *Ccl22*, and *Il21r,* associated with an M2 phenotype (Cerezo-Wallis et al., 2020), were found to be consistently downregulated. Gene set enrichment analysis further revealed that pathways related to DNA damage, cell cycle, and growth were also downregulated (**Fig. 1F**), consistent with previous studies (Cohen-Sharir et al., 2020; Wang et al., 2021). These downregulated pathways align with CIN-associated checkpoints on cell growth.

Downregulation of M2 macrophage markers and cytokines in CIN-afflicted tumors *in vivo* led us to address whether CIN in cancer cells might influence macrophages via the secretome, as suggested from previous *in vitro* studies of bone marrow-derived macrophages (BMDMs) (Cerezo-Wallis et al., 2020; Xian et al., 2021). Conditioned media from MPS1i-treated or DMSO-treated B16F10 cells was collected and added for 24 h to differentiated BMDMs. BMDMs were then processed for single-cell RNA-sequencing (**Fig. 1G**), and dimensionality reduction of gene expression via uniform manifold approximation and projection (UMAP) identified four distinct macrophage population clusters. Two clusters (0 and 2) consisted of ∼75% of macrophages treated with MPS1i-treated B16F10 secretome, and two clusters (1 and 3) consisted of ∼75% of macrophages treated with DMSO-treated B16F10 secretome (**Fig. 1H**). The latter showed increased expression of M2-like pro-cancer and anti-inflammatory markers (**Fig. 1I**). Consistent with the whole-tumor bulk RNA-sequencing, expression was increased for *Mrc1* in both clusters 1 and 3 and for *Ccl2*, *Ccl4*, and *Ccl7* in cluster 3. More importantly though, these same clusters 1 and 3 tended to downregulate M1-like anti-cancer and pro-inflammatory genes (**Fig. 1J**). Clusters 0 and 2 that consisted mostly of MPS1i-treated B16F10 secretome showed increased expression of pro-inflammatory, M1-like anti-cancer markers and little-to-no expression of most M2-like, pro-cancer markers (**Fig. 1I,J**). These results suggest that early-stage CIN in cancer cells activate macrophages via secreted factors toward M1-like anti-cancer activity.

### CIN-afflicted, CD47 knockout tumoroids are eliminated by macrophages

To assess functional effects of macrophage polarization, we focused on a 3D “immuno-tumoroid” model in which macrophage activity can work over many days against a solid proliferating mass of cancer cells in non-adherent round bottom wells (**Fig. 2A**) (Dooling et al., 2023). We chose this model primarily because 2D phagocytosis assays fail to consider the mechanical cohesive forces between tumor cells and other biophysical aspects that can affect immune response (Nia et al., 2020). We used CD47 knockout (KO) B16F10 cells, which removes the inhibitory effect of CD47 on phagocytosis, noting that KO does not perturb surface levels of Tyrp1, which is targetable for opsonization with anti-Tyrp1 (**Fig. S2A**). BMDMs were added to pre-assembled tumoroids at a 3:1 ratio, and we first assessed surface protein expression of macrophage polarization markers. Consistent with our whole-tumor bulk RNA-sequencing and also single-cell RNA-sequencing of BMDM monocultures (**Fig. 1E, 1I-J**), BMDMs from immune-tumoroids of MPS1i-treated B16F10 showed increased surface expression of M1-like markers MHCII and CD86 while showing decreased expression of M2-like markers CD163 and CD206 (**Fig. 2B-C**). Although these macrophages seemed poised for anti-cancer activity, the cancer cells showed decreased binding of anti-Tyrp1 (**Fig. S2B**) and ∼20% larger size in flow cytometry (**Fig. S2C**). The latter likely reflects cytokinesis defects and polyploidy as acute effects of CIN induction (Chunduri & Storchová, 2019; Mallin et al., 2022). Such cancer cell changes might explain why standard 2D phagocytosis assays show BMDMs attached to rigid plastic engulf relatively few anti-Tyrp1-opsonized cancer cells pretreated with MPS1i versus DMSO (**Fig. S2D**). In such cultures, BMDMs use their cytoskeleton to attach and spread, competing with engulfment of large and poorly opsonized targets. Noting that tumors *in vivo* are not as rigid as plastic, our 3D immune-tumoroids eliminate attachment to plastic, and large numbers of macrophages can cluster and cooperate in engulfing cancer cells in a cohesive mass (Dooling et al., 2023). We indeed find CIN-afflicted tumoroids are eliminated by BMDMs regardless of anti-Tyrp1 opsonization (**Fig. 2D-E**), whereas anti-Tyrp1 is required for clearance of DMSO control tumoroids (**Fig. 2D, S3B**). Imaging also suggests that cancer CIN stimulates macrophages to cluster (compare Day 4 in **Fig. 2D**), which favors cooperative phagocytosis of tumoroids (Dooling et al., 2023), and occurs despite the lack of cancer cell opsonization and their larger cell size. The 3D immune-tumoroid results with induced CIN are thus consistent with a more pro-phagocytic M1-type polarization (**Fig.1J and 2B,C**).

**Figure 2.**
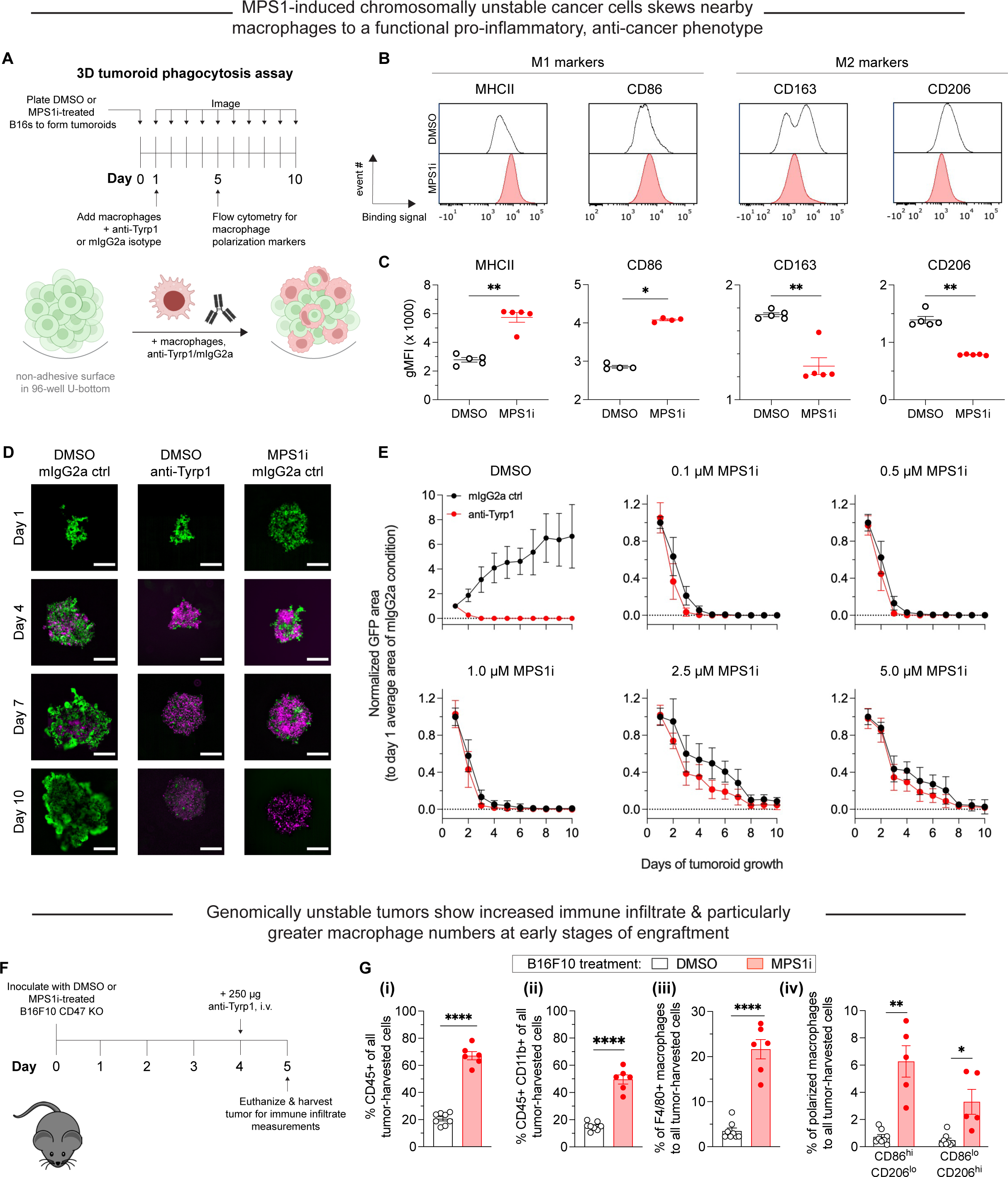
Chromosomally unstable tumoroids and tumors in early stages show increased anti-cancer macrophage polarization and activity. **(A)** Timeline and schematic for generating engineered “immuno-tumoroids” for time-lapsed studies of macrophage-mediated phagocytosis of cancer cells. Tumoroids are formed by plating and culturing B16F10 cells on non-adhesive surfaces in U-bottom shaped wells. Prior to plating for tumoroid formation, B16F10 were treated with either DMSO or MPS1i (reversine) as depicted in Fig. 1A. ∼24 h after plating, bone marrow-derived macrophages (BMDMs) with or without opsonizing anti-Tyrp1 are added to the cohesive B16F10 tumoroids. Immuno-tumoroids are imaged at the listed timepoints. At day 5, a separate subset of prepared immuno-tumoroids were collected, dissociated, and stained for measurement of macrophage polarization markers by flow cytometry. **(B)** Representative flow cytometry histograms for flow cytometry measurements of macrophage polarization markers. For M1-like (anti-tumor) markers, we chose MHCII and CD86. For M2-like (pro-tumor) markers, we chose CD163 and CD206. **(C)** Geometric mean fluorescence intensity quantification by flow cytometry of anti-tumor macrophage markers (MHCII and CD86) and pro-tumor macrophage markers (CD163 and CD206) in immuno-tumoroid cultures, with either DMSO or MPS1i-treated B16F10 CD47 KO cells. Generally, immuno-tumoroid cultures with MPS1i-treated B16F10 CD47 KO cells induce surface expression of M1-like, anti-tumor markers in macrophage while also reducing expression of M2-like, pro-tumoral markers. This suggests that, in addition to reduced cancer cell proliferation, chromosomal instability in cancer cells in early stages can prime a microenvironment conducive to anti-tumor macrophages. All data were collected from five independent immune-tumoroid culture experiments (one 96-well per replicate). Statistical significance was calculated by an unpaired two-tailed t-test with Welch’s correction (** p < 0.01). **(D)** Representative fluorescence images depicting either growth or repression of B16F10 CD47 KO cells (green) in immuno-tumoroids from days 1-5. BMDMs, shown in magenta, were added at a 3:1 ratio to initial B16F10 numbers after the day 1 images were acquired. Scale bar = 0.5 mm. **(E)** Tumoroid growth was measured by calculating the GFP+ area at the indicated timepoints (mean *±* SD, n = 16 total tumoroids from two independent experiments for each condition). All data were then normalized to average GFP+ area on day 1 of each drug treatment’s respective mouse IgG2a isotype control condition. Overall, macrophages can clear MPS1i-treated B16F10 cells regardless of either MPS1i treatment concentration or IgG opsonization. **(F)** Experimental timeline for immune cell infiltration analyses in tumors comprised of either DMSO or MPS1i-treated B16F10 CD47 KO cells. Prior to tumor inoculation, B16F10 CD47 KO cells were treated with 2.5 μM MPS1i (reversine) or the equivalent volume of DMSO vehicle control. Cells were treated for 24 h, after which they were washed twice with PBS and allowed to recover for an additional 48 h. After the recovery period elapsed, all C57BL/6 mice were subcutaneously injected with 2x10^5^ B16F10 cells. Four days (96 h) post-challenge, mice were treated with 250 μg of anti-Tyrp1 or mouse IgG2a isotype. Mice were then euthanized 24 h after antibody treatment, and their tumors were excised and disaggregated for immune infiltrate analysis by flow cytometry. **(G)** Immune cell infiltrate measurements of B16F10 CD47 KO tumors comprised of DMSO or MPS1i-treated cells. **(i)** Quantification of the percentage of CD45+ (immune) cells in the excised tumors, showing that tumors comprised of MPS1i-treated cells show ∼3-fold increased tumor immune cell infiltrate compared to their DMSO counterparts. **(ii)** Quantification of the percentage of tumor infiltrating myeloid cells in the excised tumors. B16F10 CD47 KO tumors comprised of MPS1i-treated cells show ∼2.5-fold more myeloid cells compared to their DMSO counterparts. **(iii)** Quantification of tumor infiltrating F4/80+ macrophages relative to the total number of tumor cells, with the tumors comprised of MPS1i-treated cells showing a ∼6-fold increase compared to their DMSO counterparts. **(iv)** Quantification of the number of M1-like macrophages (CD86^hi^CD206^lo^) and M2-like macrophages (CD86^lo^CD206^hi^) relative to the total number of macrophages in each tumor condition. Tumors comprised of MPS1i-treated cells have ∼6-fold more CD86^hi^CD206^lo^ compared to their DMSO counterparts. They also have a similar increase in CD86^lo^CD206^hi^ macrophages, but the number of CD86^hi^CD206^lo^ macrophages was still overall twice as high. Mean *±* SEM shown (n = 8 mice challenged with DMSO-treated B16F10 cells, n = 5 mice challenged with MPS1i-treated B16F10 cells). Statistical significance was calculated by an unpaired two-tailed t-test with Welch’s correction (ns, not significant; ** p< 0.01; **** p < 0.0001).

Given that our 3D immune-tumoroids suggest cancer cell CIN can induce anti-cancer macrophage phenotypes to favor macrophage-mediated clearance, two other MPS1i drugs were similarly tested. Tumoroids composed of AZ3146-treated cells show that BMDMs could suppress growth at high drug concentrations (**Fig. S3C-i**), whereas tumoroids composed of BAY12-17389-treated cells give results similar to reversine results (**Fig. S3C-ii**). AZ3146-treated cells tended to proliferate at most drug doses, whereas at least low doses of BAY 12-17389 and reversine also allowed for tumoroid growth (**Fig. S3D**). The results nonetheless suggest that the anti-cancer macrophage activity observed may require CIN to be severe enough to impede proliferation. Otherwise, low-level CIN may be tolerable, eventually favoring evolution (Vasudevan et al., 2020), and allowing cancer cells to proliferate more rapidly than the kinetics of phagocytosis.

Lastly, prior to long-term *in vivo* studies, we sought to expand on some of our short-term results in our analyses of transcriptomes and tumoroid surface markers. Tumors were established in mice with either MPS1i-treated or DMSO-treated B16F10 cells (per **Fig. 1D**), treated at day-4 with antiTyrp1, and isolated at day-5 for immune infiltrate analyses (**Fig. 2F, S5**). Flow cytometry showed the CIN-afflicted tumors had 3-fold more CD45+ immune cells (**Fig. 2G-i**), 2.5-fold more CD45+ CD11b+ (myeloid) cells (**Fig. 2G-ii**), and a ∼6-fold increase in F4/80-positive macrophage (**Fig. 2G-iii**) relative to DMSO controls. Furthermore, we found that CIN-afflicted tumors showed approximately 6-fold more CD86^hi^ CD206^lo^ (M1-like, anti-cancer) macrophages compared to their DMSO counterparts. We should note that these tumors also saw an increase in CD86^lo^ CD206^hi^ (M2-like, pro-cancer) macrophages compared to DMSO control (**Fig. 2G-iv**), but this increase is expected given that there are generally more macrophages infiltrating these tumors. Despite the increase in M2-like macrophages, these tumors made of MPS1i-treated cells still have ∼2-fold more M1-like macrophages (**Fig. 2G-iv**). These results suggest that although other immune cell types infiltrate tumors, which is consistent with upregulated surface expression of H2-K^b^ (**Fig. S4**), macrophage functional activity in vivo can again be expected to be positively affected by CIN-afflicted cancer cells.

### MPS1i-induced CIN favors tumor rejection with IgG opsonization and CD47 disruption

While previous studies show that complete CD47 ablation can favor suppression and even complete rejection of IgG-opsonized B16F10 tumors (Andrechak et al., 2022; Dooling et al., 2023; Hayes et al., 2023; Kamber et al., 2021), inter-experimental variation is high, particularly regarding complete tumor rejection and clearance. Furthermore, how CIN could influence these results is unknown. Therefore, we sought to assess if CIN in B16F10, which thus far has shown to skew macrophages toward an anti-cancer phenotype and away from pro-cancer one, can translate to improved efficacy and consistency in macrophage-oriented therapies. For these subsequent *in vivo* experiments, we first pre-treated B16F10 cells with either MPS1i or DMSO for 24 h, washed the drug out, and then allowed the cells to recover for an additional 48 h (**Fig. 3A**). After the recovery period elapsed, we established tumors in mice, with either MPS1i-treated or DMSO-treated B16F10 cells. To further test for CD47-mediated effects, we used either B16F10 CD47 KO or B16F10 sgCtrl, expressing wild-type CD47 (WT) levels. Starting four days post-challenge, mice received either anti-Tyrp1 or mIgG2a isotype control for opsonization.

**Figure 3.**
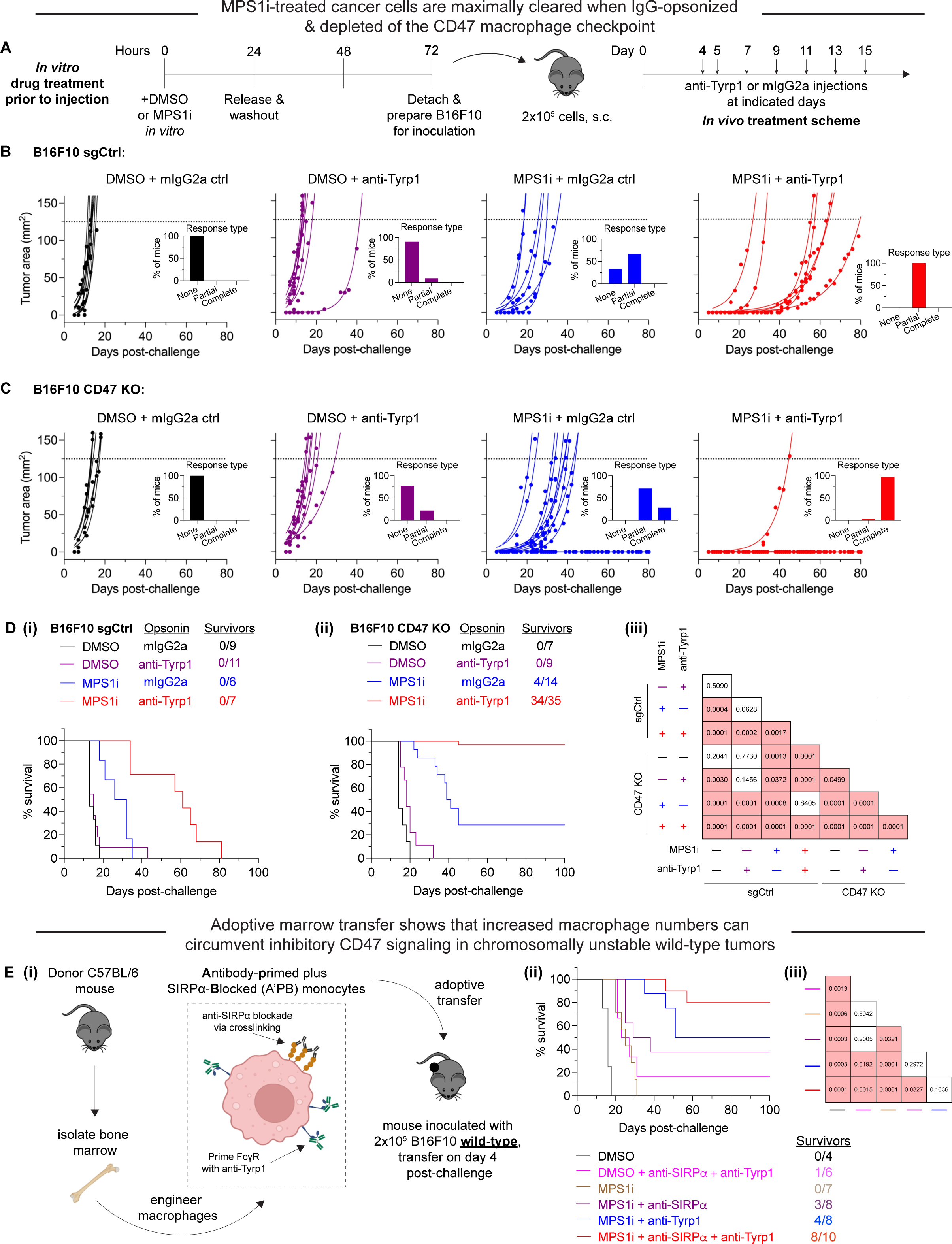
MPS1i-induced chromosomally unstable cancer cells are maximally cleared when IgG-opsonized and depleted of the CD47 macrophage checkpoint. **(A)** Timeline for *in vitro* treatment B16F10 cells prior to injection in mice and then subsequent therapeutic treatment for tumor-challenged mice. Prior to tumor inoculation, B16F10 CD47 KO cells were treated with 2.5 μM MPS1i (reversine) or the equivalent volume of DMSO vehicle control. Cells were treated for 24 h, after which they were washed twice with PBS and allowed to recover for an additional 48 h. After the recovery period elapsed, all mice were subcutaneously injected with 2x10^5^ B16F10 cells. For anti-Tyrp1 and mouse IgG2a isotype control treatments, mice were treated intravenously or intraperitoneally with 250 μg with antibody on days 4, 5, 7, 9, 11, 13, and 15 after tumor challenge. **(B)** Tumor growth curve of projected tumor area versus days after tumor challenge, with B16F10 sgCtrl cells (expressing WT levels of CD47). Each line represents a separate tumor and is fit with an exponential growth equation: A = A_0_e^kt^. Experimental conditions are as follows: n = 9 mice that were challenged with DMSO-treated B16F10 sgCtrl and subsequently treated with mouse IgG2a control, n = 11 mice that were challenged with DMSO-treated B16F10 sgCtrl and subsequently treated with anti-Tyrp1, n = 6 mice that were challenge with MPS1i-treated B16F10 sgCtrl and subsequently treated with mouse IgG2a control, and n = 7 mice that were challenge with MPS1i-treated B16F10 sgCtrl and subsequently treated with anti-Tyrp1. All data were collected across three independent experiments. Inset bar graphs depict response type for each indicated tumor challenge. A partial response was defined as a mouse that survived at least one week (20+ days) beyond the median survival of the B16F10 sgCtrl cohort treated with mouse IgG2a isotype control (13 days). **(C)** Tumor growth curve of projected tumor area versus days after tumor challenge, with B16F10 CD47 KO cells. Each line represents a separate tumor and is fit with an exponential growth equation: A = A_0_e^kt^. Complete anti-tumor responses in which a tumor never grew are depicted with the same symbol as their growing counterparts and with solid lines at A = 0. Experimental conditions are as follows: n = 7 mice that were challenged with DMSO-treated B16F10 CD47 KO and subsequently treated with mouse IgG2a control, n = 9 mice that were challenged with DMSO-treated B16F10 CD47 KO and subsequently treated with anti-Tyrp1, n = 14 mice that were challenge with MPS1i-treated B16F10 CD47 KO and subsequently treated with mouse IgG2a control, and n = 35 mice that were challenge with MPS1i-treated B16F10 CD47 KO and subsequently treated with anti-Tyrp1. All data in which mice were challenged with DMSO-treated cells were collected from three independent experiments. Data for the condition in which mice were challenged with MPS1i-challenged cells and then given mouse IgG2a control were collected from four independent experiments. Data for the final condition in which mice were injected with MPS1i-treated cells and then treated with anti-Tyrp1 was collected from seven independent experiments. Inset bar graphs depict response type for each indicated tumor challenge. A partial response was defined as a mouse that survived at least one week (20+ days) beyond the median survival of the B16F10 sgCtrl cohort treated with mouse IgG2a isotype control (13 days). **(D)** Survival curves of mice up to 100 days after the tumor challenges in (B) and (C). **(i)** Survival curves for mice challenged with B16F10 sgCtrl. **(ii)** Survival curves for mice challenged with B16F10 CD47 KO. **(iii)** Triangular matrix depicting p-values between the different tested *in vivo* conditions. Statistical significance was determined by the Log-rank (Mantel-Cox) test. **(E) (i)** Schematic depicting the different engineering anti-tumor macrophage strategies that can be used for validating macrophages’ role in clearing CIN-afflicted B16F10 cells. Fresh bone marrow was isolated from the tibia of donor C57BL/6 mice and was then incubated with anti-SIRPα clone P84 (18 μg/mL) or anti-Tyrp1 (1 μg/mL) only, both together added subsequently, or neither. Donor marrow (2x10^7^ cells) is then injected intravenously into C57BL/6 harboring B16F10 WT tumors on day 4 post-challenge. Additional anti-Tyrp1 injections, when necessary, are done on days 5, 7, 9, 11, 13, and 15 post-tumor challenge. **(ii)** Survival curves of mice up to 100 days after the B16F10 WT tumor challenge. All mice were initially challenged with 2x10^5^ B16F10 WT cells. **(iii)** Triangular matrix depicting p-values between the different tested *in vivo* conditions. Statistical significance was determined by the Log-rank (Mantel-Cox) test.

As expected, all tumors comprised of DMSO-treated B16F10 sgCtrl showed exponential growth and no survivors (**Fig. 3B**), consistent with previous studies (Dooling et al., 2023; Hayes et al., 2023). These results re-confirm the inhibitory effect that CD47 has on macrophage-mediated immunity (Ingram et al., 2017; Sockolosky et al., 2016; Willingham et al., 2012), suppressing macrophage immune response even with anti-Tyrp1 IgG opsonization. Mice with CIN-afflicted B16F10 sgCtrl tumors ultimately showed exponential growth and no survivors as well, but they did show increased median survival. Furthermore, median survival increased even further when mice were treated with anti-Tyrp1, such that all mice were considered partial responders (survival of 20+ days, one week longer than median survival of tumors comprised of DMSO-treated B16F10 sgCtrl without anti-Tyrp1 treatment). All tumors comprised of DMSO-treated B16F10 CD47 KO showed exponential growth and no survivors (**Fig. 3C**), regardless of anti-Tyrp1 treatment or not. However, CIN-afflicted B16F10 CD47 KO tumors showed more positive outcomes. Even without anti-Tyrp1, all challenged mice with CIN-afflicted CD47 KO tumors were either cured completely (28%) or considered partial responders. When paired with anti-Tyrp1 treatment, 97% of mice challenged with CIN-afflicted CD47 KO tumors survive.

Long-term survival results show that challenging mice with both MPS1i-treated and DMSO-treated B16F10 sgCtrl failed to yield any survivors, regardless of anti-Tyrp1 opsonization or not (**Fig. 3D-i**). Similarly, we also failed to generate any survivors among mice challenged with DMSO-treated B16F10 CD47 KO cells, regardless of anti-Tyrp1 opsonization or not (**Fig. 3D-ii**). This again highlights a challenge in optimizing macrophage-mediated therapies: achieving consistency in long-term therapeutic outcomes. Mice challenged with MPS1i-treated B16F10 CD47 KO cells, however, were able to survive, both without anti-Tyrp1 (28% survival) and with anti-Tyrp1 (97% survival) (**Fig. 3D-ii**). To determine if the degree of CIN affected survival, we also challenged mice with B16F10 CD47 KO cells treated with varying concentrations of reversine. Regardless of MPS1i concentration, >80% of mice survive when also treated with anti-Tyrp1 (**Fig. S6**). These results show that, in physiologically relevant microenvironments, early-stage CIN can favor survival when paired with IgG opsonization and CD47 disruption. This further suggests that macrophages are key effector cells in achieving survival against CIN-afflicted tumors, since 97% survival was achieved under conditions of maximal phagocytosis.

Although no mice challenged with CIN-afflicted B16F10 sgCtrl survived, MPS1i increase median survival significantly when paired with anti-Tyrp1 opsonization (**Fig. 3D-i**). This suggests that macrophages still display some anti-cancer activity, despite the inhibitory effects of CD47, and are important effector cells in final therapeutic outcome. To better support the hypothesis that macrophages are indeed key effector cells in rejecting CIN-afflicted tumors, we established tumors comprised of wild-type (WT) B16F10 in mice. Although WT tumors are generally unaffected by anti-CD47 and anti-Tyrp1 (Dooling et al., 2023; Hayes et al., 2023), we hypothesized that we could eliminate CIN-afflicted WT B16F10 tumors by providing adoptive transfer of marrow to increase macrophage numbers to compensate for the CD47-mediated inhibition of endogenous macrophages. Furthermore, we engineered marrow by priming Fcy receptors with anti-Tyrp1, initially inhibiting CD47-SIRPα interaction via anti-SIRPα antibody blockade or combining both (**Fig. 3E-i**). As expected, mice challenged with CIN-afflicted WT tumors and treated with regular, unprimed marrow only had ∼14% survival (**Fig. 3E-ii & iii**). This result, however, was identical to chromosomally stable WT tumors treated with marrow primed with both anti-SIRPα and anti-Tyrp1. These results suggest increased macrophages numbers do provide some minor benefit. As the marrow engineering becomes more rigorous to maximize phagocytosis, survival improves. 37% of mice challenged with CIN-afflicted WT tumors survived when treated with marrow that was initially engineered with anti-SIRPα. 50% survived with marrow primed with anti-Tyrp1. Lastly, 80% survived with anti-Tyrp1-primed and anti-SIRPα-blocked monocytes (**Fig. 3E-ii & iii**). Altogether, these results support three conclusions: (1) the increase in survival attributed to anti-SIRPα supports the idea that CD47 modulates therapeutic outcome even during CIN; (2) the increase in survival attributed to anti-Tyrp1 highlights the importance of IgG opsonization; (3) the high success from the combination emphasizes macrophages playing a key role in achieving complete rejection and clearance.

### Clearance of CIN-afflicted tumors promotes *de novo* pro-phagocytic & anti-cancer IgG

Macrophages and related phagocytic myeloid cells constitute a first line of innate immune defense against pathogens and disease, but some can also initiate acquired immunity. Two key branches of such immunity are humoral immunity, mediated by macromolecules such as antibodies, and cell-mediated immunity, involving T cells, for example. We hypothesized that mice that survived challenges with CIN-afflicted tumors would show signs of acquired immunity due to their high survival rate, and we focused on IgG because of our many functional assays.

We collected convalescent serum from survivors to quantify *de novo* anti-cancer IgG antibodies that possibly resulted from successful rejection and clearance of CIN-afflicted tumors (**Fig. 4A**). Sera was collected and then subsequently used in antibody binding testing and Western blotting to confirm emergence of anti-B16F10 antibodies. We first quantified IgG2a and IgG2b titers in convalescent sera, both of which have been previously found to engage mouse macrophage Fcy receptors (Bruhns, 2012; Nimmerjahn et al., 2010) that are typically required for macrophage-mediated phagocytosis. B16F10 cells, either Tyrp1-expresing or Tyrp1 KO, were incubated with sera (convalescent from survivors or naïve from unchallenged mice) and then counterstained with conjugated antibodies against IgG2a/c and IgG2b (**Fig. 4B-i & ii**). All mice that survived challenges from CIN-afflicted tumors yielded sera that showed significantly large increases in IgG2a/c binding against both Tyrp1-positive and Tyrp1 KO B16F10 cells. We similarly saw a statistically significant increases in IgG2b binding against both Tyrp1-positive and Tyrp1 KO B16F10 cells using the same sera, although the increases in binding were more variable. The increases in binding observed against Tyrp1 KO cells also suggest that IgG antibodies target a repertoire of antigens beyond Tyrp1, which we further qualitatively confirmed via Western blotting (**Fig. 4B-iii**). Convalescent sera were used to immunoblot against B16F10 lysates, revealing many bands at multiple molecular weights and more bands than when immunoblotting with naïve sera, supporting our hypothesis of antigen broadening beyond Tyrp1. Lastly, we also tested IgG2a and IgG2b titers from additional *in vivo* experiments: survivors from both titrated CIN-afflicted tumors from **Fig. S6** and adoptive marrow transfers in **Fig. 3E**. Similarly, convalescent sera from all these mice show increases in IgG2a/c and IgG2b in both Tyrp1-expressing and Tyrp1 KO B16F10 cells (**Fig. S7**). These results confirm that regardless of the method used to exploit CIN, induction of anti-cancer IgG can be expected.

**Figure 4.**
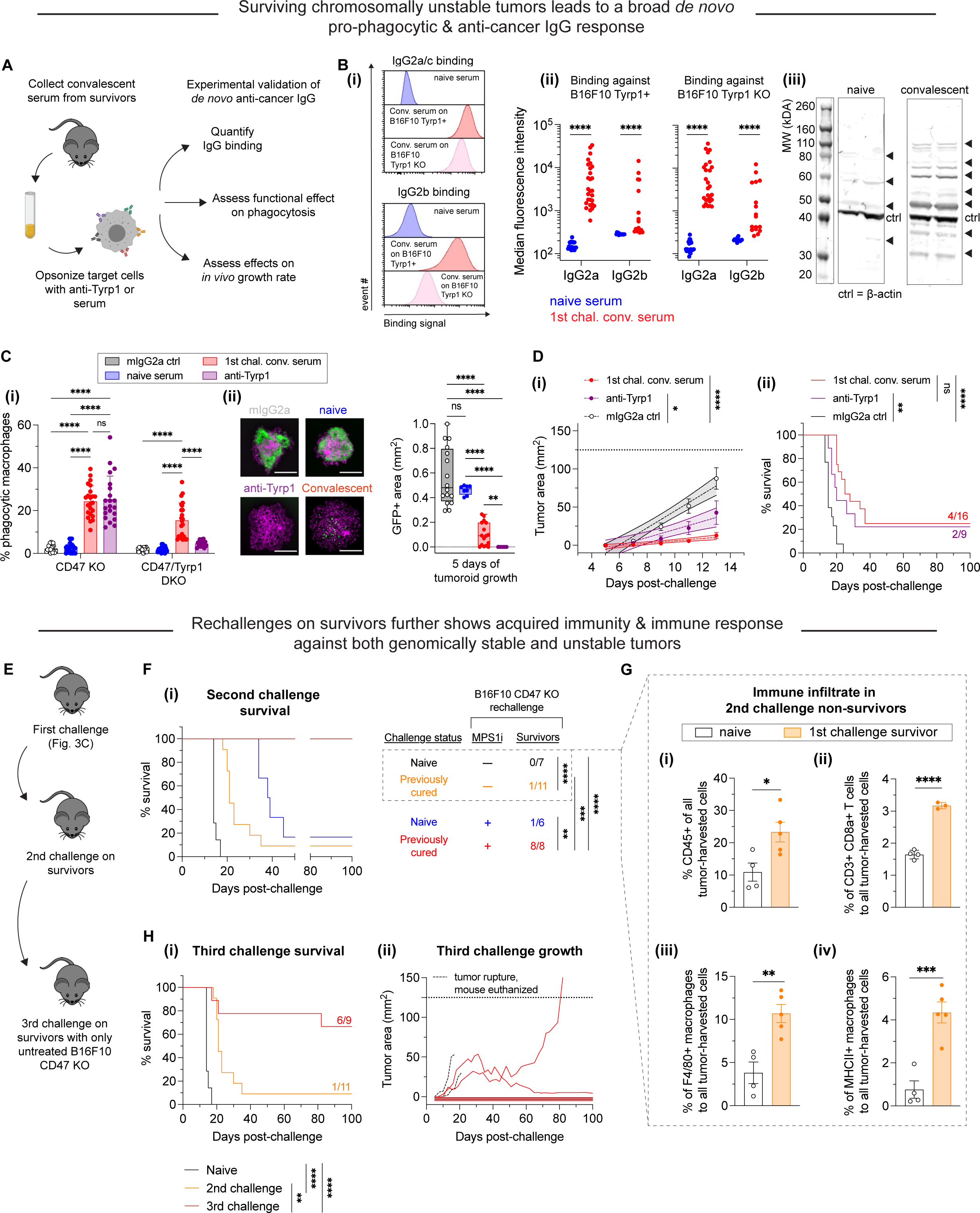
MPS1i-induced chromosomal instability favors induction of pro-phagocytic *de novo* IgG and can lead to durable acquired immunity. **(A)** Schematic illustrating protocol for sera collection from surviving mice from Fig. 3C-D and follow-up experiments to characterize potential *de novo* anti-cancer IgG antibodies and their functionality both *in vitro* and *in vivo*. Serum from all mice was collected at least 100 days after initial tumor challenge. **(B) (i)** Representative flow cytometry histograms showing that convalescent sera from survivors in Fig. 3C-D contain IgG2a/c (top) and IgG2b (bottom) that bind to both WT and Tyrp1 KO B16F10 cells. **(ii)** Median fluorescence intensity quantification of IgG2a/c and IgG2b binding from sera from surviving mice. Convalescent sera show statistically significant increase in both IgG2a/c and IgG2b titers. Binding to even Tyrp1 KO cells suggests broader recognition of antigens unique to B16F10. Statistical significance was calculated by an unpaired two-sample Kolmogorov-Smirnov test (**** p < 0.0001). For IgG2a/c quantification: for binding against Tyrp1+ cells, n = 23 distinct naïve serum samples and n = 28 distinct convalescent serum samples from surviving mice; for binding against Tyrp1 KO cells, n = 23 distinct naïve serum samples and n = 27 distinct convalescent serum samples. For IgG2b quantification: for binding against Tyrp1+ cells, n = 14 distinct naïve serum samples and n = 16 distinct convalescent serum samples from surviving mice; for binding against Tyrp1 KO cells, n = 13 distinct naïve serum samples and n = 17 distinct convalescent serum samples. **(iii)** Western blotting of B16F10 lysate with either naïve sera or first challenge survivor sera as primary probe followed by anti-mouse IgG [H+L] secondary staining. Numerous bands appear when immunoblotting with convalescent survivor sera (and more than when immunoblotting with naïve sera), qualitatively confirming binding to numerous antigens and suggesting acquired immunity beyond Tyrp1. **(C) (i)** Phagocytosis of serum-opsonized CD47 KO or CD47/Tyrp1 double KO B16F10 cells by BMDMs on 2D tissue culture plastic. Additionally, B16F10 cells opsonized with either anti-Tyrp1 or mouse IgG2a were included as controls for comparisons. Serum IgG derived from survivors has both opsonization and pro-phagocytic functional ability against B16F10. Furthermore, convalescent sera IgG from survivors is still able to drive engulfment of Tyrp1 KO cells, further suggesting targeting of antigens beyond Tyrp1. Statistical significance was calculated by two-way ANOVA and Tukey’s multiple comparison test (mean *±* SD, n = 21-23 distinct sera samples collected from survivors for B16F10 CD47 KO phagocytosis per condition and n = 14-23 for B16F10 CD47/Tyrp1 double KO phagocytosis per condition). **(ii)** Convalescent sera from first challenge survivors can repress growth of B16F10 CD47 KO immuno-tumoroids (with macrophages). Tumoroid growth was measured by calculating the GFP+ area at the indicated timepoints (mean *±* SD, n = 16 total tumoroids from two independent experiments for each condition, except n = 8 for opsonization with sera from naïve mice). Statistical significance was calculated by Brown-Forsythe and Welch ANOVA tests with Dunnett T3 corrections for multiple comparisons (ns, not significant; ** p < 0.01; **** p < 0.0001). Scale bars = 0.5 mm. **(D)** B16F10 CD47 KO cells were pre-opsonized with either convalescent sera from first challenge survivors, anti-Tyrp1 or mouse IgG2a isotype control. All mice were subcutaneously injected with 2x10^5^ pre-opsonized B16F10 CD47 KO cells. **(i)** Tumor growth curves shown are at early timepoints where growth is still in the linear regime. Linear fits highlight growth suppression of tumors comprised of B16F10 CD47 KO cells pre-opsonized with convalescent sera and anti-Tyrp1, compared to the mouse IgG2a isotype counterparts. Mean *±* SEM for all timepoints, with n = 16 mice with tumors pre-opsonized with convalescent sera (each from a distinct survivor), n = 9 mice with tumors pre-opsonized with anti-Tyrp1, and n = 14 mice with tumors pre-opsonized with mouse IgG2a isotype control. Statistical significance was calculated by ordinary one-way ANOVA and Tukey’s multiple comparison test at days 9, 11, and 13 (* p < 0.05; **** p < 0.0001). Significance represented in plot legend is representative of all three timepoints. **(ii)** Survival curves up to 100 days of mice from (D-i) with pre-opsonized tumors. Both convalescent sera and anti-Tyrp1 provide similar survival benefits, suggesting potent *de novo* IgG opsonization and anti-cancer function. Statistical significance was determined by the Log-rank (Mantel-Cox) test (ns, not significant; ** p < 0.01; **** p < 0.0001). **(E)** Schematic illustrating the series of experimental tumor challenges to assess acquired immunity. Survivors from the first challenge (Fig. 3C-D) were again challenged with either DMSO or MPS1i-treated B16F10 CD47 KO cells. Survivors from this second tumor challenge were once again challenged, this time with untreated B16F10 CD47 KO. **(F)** Survival curves of survivors from Fig. 3C-D for a second tumor challenge experiment. Prior to tumor inoculation, B16F10 CD47 KO cells were treated with 2.5 μM MPS1i (reversine) or the equivalent volume of DMSO vehicle control. Cells were treated for 24 h, after which they were washed twice with PBS and allowed to recover for an additional 48 h. After the recovery period elapsed, all mice were subcutaneously injected with 2x10^5^ B16F10 CD47 KO cells. Experimental conditions are as follows: n = 7 age-matched naïve mice (never tumor-challenged) injected with DMSO-treated B16F10 CD47 KO cells, n = 11 surviving mice (from Fig. 3C-D) injected DMSO-treated B16F10 CD47 KO cells, n = 6 age-matched naïve mice injected with MPS1i-treated B16F10 CD47 KO cells, and n = 8 surviving mice (from Fig. 3C-D) injected with MPS1i-treated B16F10 CD47 KO cells. Previous survivors challenged with DMSO-treated B16F10 CD47 KO cells show increased median survival (21 days) compared to their naïve counterpart (14 days). All previous survivors that were again challenged with MPS1i-treated B16F10 CD47 KO cells survive. All mice challenged were from three independent experiments. Statistical significance was determined by the Log-rank (Mantel-Cox) test (** p < 0.01; *** p < 0.001; **** p < 0.0001). **(G)** Non-survivors from the second tumor challenge in (F) were euthanized after tumor size was >150 mm^2^, and their tumors were excised and disaggregated for immune infiltrate analysis by flow cytometry. **(i)** Quantification of CD45+ (immune) cells in the excised tumors, showing that first challenge survivors still show ∼2.5-fold increased tumor immune cell infiltrate despite reaching terminal burden. n = 4 mice for age-matched naïve control, n = 5 mice that survived the first tumor challenge. **(ii)** Quantification of tumor infiltrating CD8a+ cytotoxic T cell relative to the total number of tumor cells. First challenge survivors show ∼2-fold increase in CD8a+ T cells. n = 4 mice for age-matched naïve control, n = 3 mice that survived the first tumor challenge. **(iii)** Quantification of tumor infiltrating F4/80+ macrophages relative to the total number of tumor cells. First challenge survivors show ∼3-fold increase in macrophages. n = 4 mice for age-matched naïve control, n = 5 mice that survived the first tumor challenge. **(iv)** Quantification of MHCII+ tumor infiltrating F4/80+ macrophages relative to the total number of F4/80 macrophages. First challenge survivors show ∼3-fold increase in MHCII+ macrophages. n = 4 mice for age-matched naïve control, n = 5 mice that survived the first tumor challenge. For all experiments, mean *±* SEM shown, and statistical significance was calculated by an unpaired two-tailed t-test with Welch’s correction (* p < 0.05; ** p < 0.01; *** p < 0.001; **** p < 0.0001). **(H) (i)** Third tumor challenge survival curves of long-term survivors from (G). All mice were challenged with 2x10^5^ B16F10 CD47 KO cells (n = 9 mice from three independent experiments), delivered subcutaneously. For benchmarking and statistical comparison, survival curves from Fig. 3F for naïve mice (n = 7 mice from three independent experiments) and second challenge (n = 11 mice from three independent experiments) are included. Long-term survivors challenged a third time show ∼70% survival without any additional therapeutic modality, suggesting significantly improved acquired immune response. Statistical significance was determined by the Log-rank (Mantel-Cox) test (** p < 0.01; *** p < 0.001; **** p < 0.0001). **(ii)** Individual tumor growth curves for third challenge in long-term survivors (n = 9 mice from three independent experiments) shown in (H-i). In total, 4 mice developed tumors, two of which had to be euthanized prematurely due to tumor rupture despite not reach a terminal burden of 125 mm^2^. These mice are still considered and included as casualties in the survival curve analysis. The two remaining mice show significantly slower tumor growth than naïve mice challenged with regular B16F10 CD47 KO (median survival of 14 days) and can be considered durable partial responders.

To test where these *de novo* serum antibodies functionally promote macrophage-mediated phagocytosis, we performed conventional 2D phagocytosis assays in which cancer cell suspensions were opsonized with sera (or anti-Tyrp1 or mouse IgG2a isotype as controls) (**Fig. 4C-i**). Under conditions of CD47 KO, we see that nearly all unpurified convalescent sera increased phagocytosis relative to naïve serum and mIgG2a isotype control. Furthermore, this increase in phagocytosis is identical to that provided by anti-Tyrp1 (both ∼5-fold higher than baseline). We also find that sera continue to promote phagocytosis even in B16F10 CD47/Tyrp1 double KO cells (∼3-fold higher than baseline). Convalescent sera are still able to increase phagocytosis against double KO cells, whereas anti-Tyrp1 expectedly does not drive phagocytosis due to lack of antigen. This again supports the hypothesis of acquired immunity with *de novo* IgG antibodies that target B16F10 antigens beyond Tyrp1.

Upon confirming the functional effect of *de novo* IgG in the convalescent sera, we then wondered if convalescent serum IgG would be able to suppress tumoroid growth, given that this model better captures both the biophysical microenvironment and proliferative capacity of tumors (Dooling et al., 2023). Indeed, we found that convalescent serum IgG added simultaneously with macrophages to B16F10 CD47 KO tumoroids led to either tumoroid elimination or significantly suppressed growth (**Fig. 4C-ii**), although the efficacy was less potent than anti-Tyrp1. This suggests that perhaps the polyclonal *de novo* IgGs here still lack the specificity benefits that accompany a monoclonal antibody such as anti-Tyrp1. Nonetheless, these results demonstrate induction of a generally potent anti-cancer antibody response to CIN-afflicted B16F10 in a CD47 KO context. Importantly, comparing these sera results for CIN-afflicted tumors to our recent studies of the same tumor model without CIN (Dooling et al., 2023; Hayes et al., 2023), we find similar levels of IgG induction (e.g. ∼100-fold above naive on average for IgG2a/c), similar increases in phagocytosis by sera opsonization (e.g. equivalent to anti-Tyrp1), and similar levels of suppressed tumoroid growth–including the variability.

We then proceeded to test the function of convalescent serum *in vivo* by opsonizing B16F10 CD47 KO cells just prior to subcutaneous implantation in naïve mice. For comparison, we also opsonized B16F10 CD47 KO cells with either anti-Tyrp1 (positive control) or mIgG2a control (negative control). We found that both convalescent sera and anti-Tyrp1 suppressed tumor growth by days 11 and 13 relative to mIgG2a control (**Fig. 4D-i**). Interestingly, we found that convalescent sera showed a trend of suppressing growth more than anti-Tyrp1, although this was not statistically significant. We continued to monitor all mice for long-term survival, and we found that this pre-opsonization with both convalescent sera and anti-Tyrp1 eliminated tumors in challenged mice with near identical cure rates (25% and 22%, respectively) (**Fig. 4D-ii**). Altogether, we found that the convalescent sera from mice originally challenged with chromosomally unstable tumors has potent anti-cancer effects *in vitro* and *in vivo*.

### Acquired immunity suppresses growth of chromosomally stable tumors and becomes more effective with ongoing challenges of CIN-afflicted tumors

The anti-cancer IgG antibody development in survivors led us to further hypothesize that we should see improved median survival and/or survival rate if surviving mice were re-challenged. We therefore challenged surviving mice with a second injection of either DMSO-treated or B16F10 CD47 KO cells (**Fig. 4E**). If additional survivors resulted from this experiment, we also intended to undergo a third challenge, akin to a prime-&-boost strategy for anti-cancer vaccination. It should be highlighted that starting from this second challenge, no mice received anti-Tyrp1. This was done to maximally challenge acquired immunity and to better simulate the possibility of recurrence post-therapy. Age-matched naïve mice receiving their first challenge responded similarly to the younger cohorts (**Fig. 3B**). Of the previously cured mice, only a single mouse (of 11 total) survived a challenge with DMSO-treated B16F10 cells (**Fig. 4F**). However, median survival increased (21 days) compared to their naïve counterparts (14 days), supporting the initial hypothesis of prolonged survival and consistent not only with past results indicating major benefits of a prime-&-boost approach with anti-Tyrp1 (Dooling et al., 2023) but also with the noted similarities in induced IgG levels. Survivors that were re-challenged again with MPS1i-treated cells, however, showed 100% survival, even in the absence of anti-Tyrp1 (**Fig. 4F**). Age-matched naïve mice receiving their first challenge of MPS1i-treated cells responded relatively similarly (a single survivor out of 6 total, 17% survival) to the younger cohort (28% survival). This complete success rate against a second challenge of CIN-afflicted B16F10 CD47 KO further supports an acquired immune response, at least against ongoing chromosomally unstable cells.

Mice that failed to reject re-challenge tumors comprised of DMSO-treated B16F10 CD47 KO in **Fig. 4F** were euthanized and had their tumors harvested to measure their immune cell infiltrate by flow cytometry (**Fig. S8**). Although these mice did not survive, we found that these previous survivors showed roughly a 2-fold increase in the number of immune cells in their tumors (**Fig. 4G-i**), a 2-fold increase in the number of CD3+ CD8+ T cells (**Fig. 4G-ii**), and a near 3-fold increase in the number of F4/80+ macrophages (**Fig. 4G-iii**) compared to their naïve counterparts. We further found that of the F4/80+ macrophage infiltrate, previous survivors had roughly four times as many MHCII-high macrophages (M1-like, anti-cancer). These immune infiltrate analyses suggest that although the acquired immune response in these previous survivors was still not potent enough to clear chromosomally stable and normally proliferating B16F10 CD47 KO tumors, it did enhance cell-mediated immunity.

Lastly, we performed a third tumor challenge on second challenge survivors with untreated B16F10 CD47 KO cells (chromosomally stable and regular proliferating). Mice were left untreated post-challenge (no anti-Tyrp1). ∼56% of these mice completely resisted tumor growth. Of the four mice that developed tumors, two had to be euthanized prematurely due to tumor rupture but were indeed showing signs of growth suppression. The last two mice consisted of two long-term partial responders: one whose tumor did not reach terminal burden until 82 days post-challenge and another who experienced stable tumor regression with almost no regrowth. Overall, the 56% survival rate in the third challenge, in the absence of anti-Tyrp1, and the partial responses observed in mice that grew tumors suggest a durable immunological response that results from encountering and clearing CIN tumors, at least in the context of CD47 disruption.

Macrophages seem to be the key *<u>initiating</u>-*effector cells, based in part on the following findings. First, macrophages with both SIRPα blockade and FcR-engaging, tumor-targeting IgG maximize survival of mice with WT B16 + Rev tumors (**Fig. 3E**) – noting that macrophages but not T cells express SIRPα and FcR’s. Despite the clear benefits of adding macrophages, to further assess whether T and B cells are key <u>initiating</u>-effector cells, new experiments were done with mice depleted of T and B cells. We compared the growth delay of MPS1i versus DMSO treatments in these mice to the delay in fully immunocompetent mice with T and B cells – with all studies done at the same time. We found that slower growth with Rev relative to DMSO was similar in mice without T and B cells when compared to immunocompetent C57 mice (**Fig.S9**). We conclude therefore that T and B cells are not key <u>initiating</u>-effector cells. At later times, B cells are likely effector cells at least in terms of making anti-tumor IgG, and T cells in tumor re-challenges are also increased in number (**Fig. 4G-ii**). We further note that in our earlier collaborative study (Harding et al., 2017) WT B16 cells were pre-treated by genome-damaging irradiation before engraftment in C57 mice, and these cells grew minimally – similar to MPS1i treatment – while untreated WT B16 cells grew normally at a contralateral site in the same mouse. Such results indicate that T and B cells in C57BL/6 mice are not sufficiently stimulated by genome-damaged B16 cells to generically impact the growth of undamaged B16 cells.

## Discussion

Macrophage-directed immunotherapies against solid tumors have the potential for maximal efficacy when at least three elements are combined: large numbers of macrophages for cooperativity, IgG opsonization that activates Fc receptors and stimulates macrophage-mediated phagocytosis, and disruption of the CD47 macrophage immune checkpoint (Dooling et al., 2023). However, even with these factors properly applied, complete tumor rejection and clearance is not guaranteed and varies greatly across *in vivo* mouse studies (Andrechak et al., 2022; Dooling et al., 2023; Hayes et al., 2023; Kamer et al., 2021). Here, we show a 97% survival rate of immunocompetent mice challenged with CIN-afflicted, syngeneic B16F10 tumors when maximizing macrophage-mediated activity. The result is notable given that B16F10 tumors are poorly immunogenic, do not respond to either anti-CD47 or anti-PD-1/PD-L1 monotherapies, and show modest and variable cure rates (∼20-40%; Dooling et al., 2023; Hayes et al., 2023) even when macrophages have been made maximally phagocytic according to notions above. We should note here that our whole-tumor RNA-sequencing data (**Fig.1E**) shows expression of PD-1 (gene *Pdcd1*) follows no consistent trend upon MPS1i treatment, and that *Pdcd1* was not detected in our single cell RNA-sequencing data for macrophage cultures (**Fig.1G**)–motivating further study.

These results suggest that CIN in early stages generates anti-cancer vulnerabilities that favor macrophage-mediated immune response, contingent on conditions of maximal phagocytosis. Immunocompetent mice consistently survive these challenges at high survival rates. These survivors also develop *de novo* anti-cancer IgG, similar to previous studies (Dooling et al., 2023; Hayes et al., 2023), that are pro-phagocytic, multi-epitope, and efficacious *in vivo*. The emergence of these IgGs could be synergistic with clinically relevant CD47 blockade treatments for solid tumors and help address concerns regarding resistance due to antigen loss (Jalil et al., 2020). Mice that are re-challenged with chromosomally stable CD47 KO tumors (and without exogenous anti-Tyrp1 opsonization) shows increased median survival and increased immune infiltrate, further supporting the hypothesis of newly generated anti-cancer acquired immunity. More interestingly, though, we see that all mice re-challenged with CIN-afflicted CD47 KO tumors survive, even in the absence of anti-Tyrp1 opsonization. A third challenge of these two-time survivors with chromosomally stable CD47 KO tumors shows >50% mice survive and improved median survival for non-survivors. These results elucidate at least two advantages that CIN can provide to better therapeutic outcomes. First, early-stage CIN facilitates survival while generating potent *de novo* IgGs that can drive positive phagocytic feedback to minimize recurrence. Second, ongoing vulnerability-inducing CIN in tumor cells can both strengthen cell-mediated acquired immunity and create an antigen reservoir for the maintenance of long-term humoral immunity.

The effects of CIN and aneuploidy in macrophages certainly require further investigation. We recently published that M1-like polarization of BMDMs with IFNγ priming is sufficient to suppress growth of B16 tumoroids with anti-Tyrp1 opsonization. This suppression occurred more rapidly than unpolarized/unprimed macrophages and much more rapidly than M2-like polarization of BMDMs with IL-4 (Extended Data Fig.5A in Dooling et al., 2023). We therefore conclude that anti-cancer polarization contributes in this assay. While the secretome from MPS1i-treated cancer cells has been found to trigger expression of *Arg1* and *Il6* (Xian et al., 2021), both of which are pro-cancer M2-like macrophage markers (Fernando et al., 2014; Mujal et al., 2022), our findings suggest that polarization is much more complex. Here, whole-tumor bulk RNA-sequencing hints at CIN-afflicted tumors having a macrophage population that is both less anti-inflammatory and M2-like than their stable counterparts. Single-cell RNA-sequencing of BMDMs treated with secretome from CIN-afflicted cells further suggests that CIN induces a microenvironment that can push macrophages to a pro-inflammatory, anti-cancer phenotype while minimizing polarization to a pro-cancer phenotype. Our transcriptomics analyses also align more with deeper investigation that suggest additional markers are required for macrophage polarization distinction (Jablonksi et al., 2015). We further confirm these findings by observing increased surface protein expression of anti-cancer M1-like macrophages *in vitro* 3D tumoroid co-cultures with CIN-afflicted cells and *in vivo* immune infiltrate experiments. Functional tests are also crucial: BMDMs show enhanced clearance of CIN-afflicted cells in 3D tumoroid phagocytosis assays. Additionally, the aforementioned study (Xian et al., 2021) only found this trend in SKOV3 cells, whereas their aneuploid fused B16 cells show decreased *Arg1* expression, suggesting possible cell-intrinsic complications. Meta-analyses of aneuploid tumor data from TCGA also suggest reduced anti-cancer macrophage activity (Davoli et al., 2017; Xian et al., 2021). Our study, however, distinctly highlights macrophage response at early timepoints of CIN, during the which cancer cells have not yet adapted to CIN- and aneuploidy-induced stresses. TCGA studies, on the other hand, are limited to much later timepoints when cells have overcome CIN- and aneuploidy-associated stresses and developed their own unique aneuploidies to drive tumor progression. Additionally, more recent studies find that aneuploidy and CIN paired with high tumor mutational burdens show increased median survival in patients (Spurr et al., 2022a; Spurr et al., 2022b), suggesting possible other complexities that favor immune infiltration and that could explain how CIN can favor macrophage-mediated immune response. Important to also consider–as shown here–are the coupled effects with cancer CIN of the macrophage checkpoint pathway CD47-SIRPα and its blockade in combination with tumor opsonization.

## Materials and Methods

### Cell culture

B16F10 cells (CRL-6475) were obtained from American Type Culture Collection (ATCC) and cultured at 37°C and 5% CO_2_ in either RPMI-1640 (Gibco 11835-030) or Dulbecco’s Modified Eagle Medium (DMEM, Gibco 10569-010) supplemented with 10% fetal bovine serum (FBS, Sigma F2442), 100 U/ mL penicillin and 100 μg/mL streptomycin (1% P/S, Gibco 15140-122). B16F10 cells were maintained in passage in RPMI-1640 but switched to DMEM at least three days prior to *in vivo* subcutaneous injections. 293T human embryonic kidney (CRL-1573) cells were also obtained from ATCC and cultured at 37°C and 5% CO_2_ in DMEM culture media supplemented with 10% FBS and 1% P/S. All cell lines were passaged every 2-3 days when a confluency of ∼80% was reached. For trypsinization, cells were washed once with Dulbecco’s phosphate-buffered saline (PBS, Gibco 14190-136) and then detached with 0.05% Trypsin (Gibco 25300-054) for 5 min at 37°C and 5% CO_2_. Trypsin was quenched with an equal volume of complete culture media.

### Lentiviral production and transduction

24 h prior to lentivirus production, 293T cells were plated at a density of 8x10^5^ cells in 2 mL of DMEM in individual wells of a 6-well plate. On the day of transfection, 1.35 μg of psPAX2, 165 ng of pCMV-VSV-G, and 1.5 μg of lentiviral plasmid were added into a microcentrifuge tube with 7.5 μL of Mirus TransIT-Lenti transfection reagent (Mirus Bio, 6604). Once all plasmids were pooled together with the transfection reagent, 300 μL of serum-free media was added. The solution was gently mixed by pipetting and then allowed to incubate for 30 min. After 30 min elapsed, the solution was gently added dropwise by pipetting to an individual 6-well containing the 293T cells plated the day prior. We note that we also use B16F10 CD47 and Tyrp1 KO lines in this study, whose preparation has been previously described (Hayes, et al., 2020). LentiV-cas9_puro and Lenti_sgRNA_EFS_GFP plasmids (Addgene #<u>108100</u> and <u>65656</u>, respectively) were gifts from Christopher Vakoc. psPAX2 was a gift from Didier Trono (Addgene plasmid #12260). pVSV-G was a gift from Bob Weinberg (Addgene plasmid #8454). The single guide RNA (sgRNA) oligonucleotides (CD47, 5′-TCCCCGTAGAGATTACAATG-3′; SIRPα, 5′-TAATTCTAAGGTCATCTGCG-3′) were designed using the Broad CRISPR algorithm. sgRNAs were cloned into the sgRNA vector using a BsmBI restriction digest.

Viral production was allowed to continue for 48 h, after which the supernatant was collected from each well and centrifuged for 5 min at 300 x *g*. The supernatant was collected and then either added directly to B16F10 cells (0.5 mL of lentivirus-containing supernatant to 2x10^4^ cells in an individual 6-well) or stored at -70°C for long-term storage. 72 h after transduction, spent media with lentivirus was aspirated, B16F10 cells were washed with PBS, and fresh media was added. For selecting successfully lentivirally transduced cells, cells were cultured in fresh media containing 1 μg/mL of puromycin (Invitrogen A1113803). Cells were kept in puromycin-containing until a non-transduced control population also treated with puromycin completely died.

### Antibodies

Antibodies used for *in vivo* treatment and blocking and for *in vitro* phagocytosis are as follows: anti-mouse/human Tyrp1 clone TA99 (BioXCell BE0151), mouse IgG2a isotype control clone C1.18.4 (BioXCell BE0085), and Ultra-LEAF anti-mouse CD172a (SIRPα) clone P84 (BioLegend 144036). Low-endotoxin and preservative-free antibody preparations were used for *in vivo* treatments and *in vitro* phagocytosis experiments. For primary antibody staining of surface proteins via flow cytometry, the following were used: anti-mouse CD47 clone MIAP301 (BioXCell BE0270) and anti-mouse/human Tyrp1 clone TA99. Secondary antibodies used for flow cytometry are as follows: Alexa Fluor 647 donkey anti-mouse IgG (ThermoFisher A-31571) and Alexa Fluor 647 goat anti-rat IgG (ThermoFisher A-21247). All secondary antibody concentrations used followed the manufacturer’s recommendations.

For immune infiltrate analysis, the following BioLegend antibodies were used: Brilliant Violet 650 anti-mouse CD45 clone 30-F11 (103151), Brilliant Violet 785 anti-mouse CD45 clone 30-F11 (103149), APC/Cy7 anti-mouse CD45 clone 30-F11 (103115), APC anti-mouse/human CD11b clone M1/70 (101212), PE/Cyanine7 anti-mouse/human CD11b clone M1/70 (101216), PE/Dazzle 594 anti-mouse Ly6G clone 1A8 (127647), PerCP anti-mouse Ly-6G clone 1A8 (127653), PE anti-mouse F4/80 clone BM8 (123110), Brilliant Violet 605 anti-mouse Ly-6C clone HK1.4 (128035), APC anti-mouse I-A/I-E clone M5/114.15.2 (107614), Pacific Blue anti-mouse CD86 clone GL-1 (105022), APC/Cy7 anti-mouse CD86 clone GL-1 (105029), APC anti-mouse CD206 (MMR) clone C068C2 (141707), Brilliant Violet 421 anti-mouse CD206 clone C068C2 (141717), PE/Cy7 anti-mouse CD163 clone S15049F (156707), APC/Cy7 anti-mouse CD3e clone 145-2C11 (100329), and Alexa Fluor 647 anti-mouse CD8a clone 53-6.7 (100727). TruStain FcX PLUS (anti-mouse CD16/32) clone S17011E (156603) was used in all immune infiltrate experiments to block Fc receptors. For immunogenicity post-MPS1i treatment, APC anti-mouse H-2kb/H-2Db clone 28-8-6 (114613) was used. For IgG titer in tumor challenge surviving mice, the following BioLegend antibodies were used: PE anti-mouse IgG2a clone RMG2a-62 (407108, known to bind IgG2c as well) and APC anti-mouse IgG2b clone RMG2b-1 (406711). Primary antibody used in Western blotting was anti-β-actin clone C4 (Santa Cruz sc 47778). Secondary antibody used in Western blotting was HRP sheep anti-mouse IgG (GE Life Sciences NA931V).

### Mice

C57BL/6 mice (Jackson Laboratory 000664) were 6-12 weeks old at the time of tumor challenges and for bone marrow harvesting, with the exception of second and third challenge experiments. NSG mice aged 6–12 weeks old were obtained from the Stem Cell & Xenograft Core at the University of Pennsylvania. Additionally, for re-challenge experiments, age-matched naïve mice were used. All experiments were performed in accordance with protocols approved by the Institutional Animal Care and Use Committee (IACUC) of the University of Pennsylvania.

### Drug treatments & micronuclei quantification

For MPS1i studies, the following chemical drugs were used: reversine (Cayman Chemical 10004412), AZ3146 (Cayman Chemical 19991), and BAY 12-17389 (Selleck Chemicals S8215). 24 h prior to treatment, B16F10 cells were in either 6-well or 12-well plates. For 6-wells, 20,000 cells were plated per well. For 12-wells, 2,000 cells were plated per well. On the day of treatment, spent media was aspirated, and fresh media supplemented with 10% FBS, 1% P/S, and either MPS1i or DMSO vehicle control was added to each well. The concentration used for each treatment is listed in the Figure legend. The volume of DMSO added was equal to the volume required for the highest MPS1i concentration for each experiment. All cells were treated for 24 h, after which drug-containing spent media was aspirated. Cells were then washed with a full volume of PBS for 5 min. PBS was aspirated, and two repeat washes were performed. Cells were then allowed to recover from MPS1i treatment for an additional 48 h, with a fresh media replacement 24 h after the initial wash.

For imaging and micronuclei quantification, cells were fixed with 4% formaldehyde for 20 min after the 48 h recovery period and later imaged on an Olympus IX inverted microscope with a 40x/0.6 NA objective. The Olympus IX microscope was equipped with a Prime sCMOS camera (Photometrics) and a pE-300 LED illuminator (CoolLED) and was controlled with MicroManager software v2. At least 200 B16F10 were imaged per individual well for micronuclei quantification.

### Bone marrow-derived macrophages (BMDMs)

Bone marrow was harvested from the femurs and tibia of donor mice, lysed with ACK buffer (Gibco A1049201) to deplete red blood cells, and then cultured on Petri culture dishes for 7 days in Iscove’s Modified Dulbecco’s Medium (IMDM, Gibco 12440-053) supplemented with 10% FBS, 1% P/S, and 20 ng/mL recombinant mouse macrophage colony-stimulating factor (M-CSF, BioLegend 576406). 72 h after initial plating, one whole volume of fresh IMDM supplemented 10% FBS, 1% P/S, and 20 ng/mL M-CSF. After 7 days of differentiation, spent media was removed, BMDMs were gently washed once with phosphate-buffered saline (PBS), and fresh IMDM supplemented with 10% FBS, 1% P/S, and 20 ng/mL M-CSF was added.

### Conditioned media treatment of BMDMs

BMDMs that had successfully undergone 7 days of differentiation in 20 ng/mL M-CSF were used. Spent media was removed, BMDMs were gently washed once with phosphate-buffered saline (PBS), and fresh IMDM supplemented 10% FBS, 1% P/S, and 20 ng/mL M-CSF was added. Then, conditioned media from B16F10 cells treated with either MPS1i or DMSO was collected. Conditioned media was centrifuged for 5 min at 300 x *g* to remove any cellular debris. The supernatant was collected and then supplemented to 5% FBS, 1% P/S, and 20 ng/mL M-CSF. One whole volume of conditioned media was added to BMDMs with fresh media. 24 h after treatment, BMDMs were detached using 0.05% Trypsin and processed for single-cell RNA-sequencing.

### *In vitro* phagocytosis

For two-dimensional phagocytosis assays, BMDMs were detached using 0.05% Trypsin and re-plated in either 6-well or 12-well plates, at a density of 1.8x10^4^ cells per cm^2^ in IMDM supplemented 10% FBS, 1% P/S, and 20 ng/mL M-CSF. After 24 h elapsed, BMDMs were labeled with 0.5 μM CellTracker DeepRed dye (Invitrogen C34565), according to the manufacturer’s protocol. Following staining, BMDMs were washed and incubated in serum-free IMDM supplemented 0.1% (w/v) BSA and 1% P/S. B16F10 cells were labeled with carboxyfluorescein diacetate succinimidyl ester (Vybrant CFDA-SE Cell Tracer, Invitrogen V12883), also according to the manufacturer’s protocol. B16F10 cells were detached and opsonized with 10 μg/mL anti-Tyrp1, with 10 μg/mL mouse IgG2a isotype control antibody, or 5% (v/v) mouse serum in 1% BSA. Opsonization was allowed for 30-45 min on ice. Opsonized B16F10 suspensions were then added to BMDMs at a ∼2:1 ratio and incubated at 37°C and 5% CO_2_ for 2 h. Non-adherent cells were removed by gently washing with PBS. For imaging, cells were fixed with 4% formaldehyde for 20 min and later imaged on an Olympus IX inverted microscope with a 40x/0.6 NA objective. The Olympus IX microscope was equipped with a Prime sCMOS camera (Photometrics) and a pE-300 LED illuminator (CoolLED) and was controlled with MicroManager software v2. At least 300 macrophages were imaged per individual well for calculation of phagocytosis.

### 3D tumoroid formation and phagocytosis

Briefly, non-TC-treated 96-well U-bottom plates were treated with 100 μL of anti-adherence rinsing solution (StemCell Technologies 07010) for 1 h. The cells were then washed with 100 μL of complete RPMI 1640 cell culture media. This generated surfaces conducive to generating tumoroids and preventing cells from adhering to the well bottom during experiments. B16F10 were detached by brief trypsinization, resuspended at a concentration of 1x10^4^ cells per mL in complete RPMI 1640 cell culture media (10% FBS, 1% P/S) with 50 μM β-mercaptoethanol (Gibco 21985023). 100 μL of this cell suspension was added to each well such that each tumoroid initially started with approximately 1x10^3^ cells. Aggregation of B16F10 cells was confirmed 24 h later by inspection under microcopy. Upon confirmation of tumoroid formation, BMDMs were labeled with 0.5 μM CellTracker DeepRed dye (Invitrogen C34565), according to the manufacturer’s protocol. BMDMs were then detached by brief trypsinization and gentle scraping and resuspended in complete RPMI 1640 cell culture media at a concentration that would allow for delivery of 3x10^3^ BMDMs to each individual tumoroid culture. The BMDM cell suspension was also supplemented with 120 ng/mL M-CSF and antibodies (either anti-Tyrp1 or mouse IgG2a isotype control) such that delivery of 20 μL of this suspension to each individual tumoroid culture result in final concentrations of 20 ng/mL M-CSF and 20 μg/mL of antibody. For tumoroid studies in which mouse convalescent serum was used, the BMDM cell suspension was supplemented with 120 ng/mL M-CSF and serum such that delivery of 20 μL of this suspension to each individual tumoroid culture resulted in final concentrations of 20 ng/mL M-CSF and a final mouse serum concentration of 1:200. Tumoroids were imaged on an Olympus IX inverted microscope with a 4x/0.6 NA objective.

For macrophage polarization tumoroid experiments (Fig. 2B-C), B16F10 tumoroids were prepared in the same manner as described above, with the exception that no anti-Tyrp1 or mIgG2a was added. After 5 days had elapsed for the experiment, tumoroids were dissociated by brief trypsinization and stained with conjugated antibodies targeting MHCII, CD86, CD163, and CD206, following the manufacturer’s protocol. Cells were then run on a BD LSRII (Benton Dickinson) flow cytometer. Data were analyzed with FCS Express 7 software (De Novo Software).

### *In vivo* tumor growth

B16F10 cells cultured in DMEM growth media were detached by brief trypsinization, washed twice with PBS, and resuspended at 2x10^6^ cells per mL. Cell suspensions were kept on ice until injection. All subcutaneous injections were performed on the right flank while mice were anesthetized under isoflurane. Fur on the injection site was wet slightly with a drop of 70% ethanol and brushed aside to better visualize the skin for injection beneath the skin of a 100 μL suspension of cancer cells. For assessing immune infiltrates in early stages of tumor engraftment, when tumors are still small, we used a relatively high number of tumor cells (500,000 cells in Fig. 1D and Fig. 2F-G) to achieve sufficient cell numbers after dissociating the tumors, particularly for the slow-growing MPS1i-treated tumors. More specifically, with dissection, collagenase treatment, passage through a filter to remove clumps, we would lose many cells, and yet needed 100,000 viable cells or more for bulk RNA-seq suspensions and for flow cytometry measurements. For all other studies, 200,000 cancer cells were injected, and for subsequent treatments, mice received either intravenous (I.V.) or intraperitoneal (I.P.) injections of anti-Tyrp1 clone TA99 or mouse IgG2a isotype control clone C1.18.4 (250 μg antibody in 100 μL PBS) on days 4, 5, 7, 9, 11, 13, and 15 post-tumor challenge. Intravenous injections were done via the lateral tail vein. Tumors were monitored by palpation and measured with digital calipers. The projected area was roughly elliptical and was calculated as A = π/4 x *L* x *W*, where *L* is the length along the longest axis and *W* is the width measured along the perpendicular axis. For our studies, a projected area of 125 mm^2^ was considered terminal burden for survival analyses. Mice were humanely euthanized following IACUC protocols if tumor size reached 2.0 cm on either axis, if tumor reached a projected area greater than 200 mm^2^ or if a tumor was ulcerated.

### Adoptive cell transfers

Fresh bone marrow was harvested as described in the **Bone marrow-derived macrophages (BMDMs)** methods section. Marrow cells were counted on a hemocytometer and resuspended to a concentration of 8x10^7^ cells per mL in in 5% (v/v) FBS/PBS. To block SIRPα, cells were then incubated with anti-SIRPα clone P84 (18 μg/mL) for 1 h at room temperature on a rotator. After the incubation period elapsed, cells were centrifuged at 300 x *g* for 5 min, washed with PBS, and then centrifuged once more at 300 x *g* for 5 min to remove any unbound anti-SIRPα. Cells were again re-suspended to a concentration of 8x10^7^ cells per mL in in 2% (v/v) FBS/PBS, with or without anti-Tyrp1 clone TA99 (1 mg/mL). Marrow cells were then injected intravenously (2x10^7^ cells in 250 μL per mouse) into tumor-bearing mice. All adoptive transfers were done four days post-challenge. Control data were adapted from (Dooling et al., 2023) to minimize mice for experiments, since the cited study establishes proper benchmarks for comparison and also finds that these control conditions minimally (if at all) improve survival.

### Serum collection & IgG titer quantification

Blood was drawn retro-orbitally from mice anesthetized under isoflurane, using heparin- or EDTA-coated microcapillary tubes. Collected blood was allowed to clot for 1 h at room temperature in a microcentrifuge tube. The serum was separated from the clot by centrifugation at 1,500 x *g* and stored at -20°C for use in flow cytometry and phagocytosis assays.

For IgG titer quantification, B16F10 cells were detached by trypsinization and incubated with 5% (v/v) mouse serum in 1% BSA. Opsonization was allowed for 30-45 min on ice. After the incubation period elapsed, cells were centrifuged at 300 x *g* for 5 min, washed once with PBS, centrifuged again at 300 x *g* for 5 min, and then resuspended in 0.1% (w/v) BSA with both PE anti-mouse IgG2a/c clone RMG2a-62 and APC anti-mouse IgG2b clone RMG2b-1 (see **Antibodies** section for more information). Anti-IgG conjugated-antibody incubation occurred for 30-45 min, after which cells were centrifuged at 300 x *g* for 5 min, washed once with PBS, centrifuged again at 300 x *g* for 5 min, and then resuspended in 5% (v/v) FBS/PBS. Cells were run on a BD LSRII (Benton Dickinson) flow cytometer.

### Western blotting

Lysate was prepared from B16F10 cells using RIPA buffer containing 1X protease inhibitor cocktail (Millipore Sigma P8340) and boiled in 1X NuPage LDS sample buffer (Invitrogen NP0007) with 2.5% β-mercaptoethanol. Proteins were separated by electrophoresis in NuPage 4-12% Bis-tris gels run with 1X MOPS buffer (Invitrogen NP0323) and transferred to an iBlot nitrocellulose membrane (Invitrogen IB301002). The membranes were blocked with 5% (w/v) non-fat milk in Tris buffered saline (TBS) plus Tween-20 (TBST) for 1 h and stained with 5% (v/v) mouse serum overnight at 4°C with agitation. The membranes were washed TBST and incubated with 1:500 secondary antibody conjugated with horseradish peroxidase (GRP) in 5% (w/v) milk in TBST for 1 h at room temperature with agitation. The membranes were then washed again three times with TBST. Membranes probed with HRP-conjugated secondary antibody were developed a 3,3’,5,5’-teramethylbenzidine (TMB) substrate (Genscript L002V or Millipore Sigma T0565). Developed membranes were scanned and then processed with ImageJ.

### Immune infiltrate analysis of tumors

For day 5 post-challenge measurements: If mice required anti-Tyrp1 treatment, mice received a single dose of intravenously delivered anti-Tyrp1 or mouse IgG2a isotype control four days (96 h) post-tumor challenge. 24 later, mice were humanely sacrificed. Otherwise, mice were sacrificed five days post-tumor challenge. For immune analysis of second challenge non-survivors: Mice were humanely euthanized when tumor burden reached >150 mm^2^.

Tumors from euthanized mice were excised and placed into 5% (v/v) FBS/PBS. Tumors were then disaggregated using Dispase (Corning 354235) supplemented with 4 mg/mL of collagenase type IV (Gibco 17104-019) and DNAse I (Millipore Sigma, 101041159001) for 30-45 min (until noticeable disaggregation) at 37°C, centrifuged for 5 min at 300 x *g*, and resuspended in 1 mL of ACK lysis buffer for 12 min at room temperature. Samples were centrifuged for 5 min at 300 x *g*, washed once with PBS, and then resuspended in 5% (w/v) BSA/PBS for 20 min. After 20 min elapsed, fluorophore-conjugated antibodies to immune markers were added to each cell suspension. The following markers were used for analysis: for macrophages, CD45, CD11b, F4/80, Ly-6C, Ly-6G, CD86, CD206, MHCII; for T cells, CD45, CD3e, CD8a. Antibody binding occurred for 30 mins while samples were kept on ice and covered from light. Samples were then centrifuged for 5 min at 300 x *g*, washed once with PBS, and resuspended in FluoroFix Buffer (BioLegend, 422101) for 1 h at room temperature prior to analysis on a BD LSRII (Benton Dickinson) flow cytometer. For day 5 post-challenge measurements, 100,000 to 200,000 live cells were collected. For *in vivo* tumor infiltrate studies in re-challenged mice, 10 million live cells were collected. Data were analyzed with FCS Express 7 software (De Novo Software).

### Bulk RNA-sequencing

RNA library was constructed using NEBNext® Ultra™ II RNA Library Prep Kit for Illumina® and NEBNext® Multiplex Oligos for Illumina (E7770S, E7335) per the manufacturer’s instructions. The library prepared was processed at the Next Generation Sequencing Core at the University of Pennsylvania (12-160, Translational Research Center) using NovaSeq 6000, 100 cycles (Illumina). The reads were aligned to mouse reference, mm10 (GENCODE vM23/Ensembl 98) using star alignment. Cell count matrix was generated and imported to RStudio for downstream analysis. Package “DESeq2” (v1.32.0) was used for normalization and differential expression analysis. Package “fgsea” (v1.18.0) was used for gene set enrichment analysis. Additional exploratory data analysis was then done using either RStudio or Python 3.8.

### Single-cell RNA-sequencing

RNA libraries were prepared using the 10X Genomics Chromium Single Cell Gene Expression kit (v3.1, single index, PN-1000128; PN-1000127; PN-1000213) per the manufacturer’s instructions. The libraries were sequenced at the Next Generation Sequencing Core using NovaSeq 6000, 100 cycles (Illumina). Raw base call (BCL) files were analyzed using CellRanger (version 5.0.1) to generate FASTQ files, and the “count” command was used to generate raw count matrices aligned to mm10 (GENCODE vM23/Ensembl 98). Cells were filtered to make sure that they expressed a minimum of 1,400 genes with less than 15 percent mitochondrial content. Data was normalized using the “LogNormalize’’ method from the Seurat package. Differential expression analysis was performed using the “FindAllMarkers’’ command, and the output was used for gene set enrichment analysis (GSEA).

### InferCNV

Count matrix of single cell RNA-sequencing results was used as input for InferCNV object construction (1.7.1) (InferCNV of the Trinity CTAT Project, see the following for more information: https://github.com/broadinstitute/inferCNV). Gene position files were created for GRCm38. Single-cell RNA-sequencing data of DMSO-treated B16F10 were used as reference for copy number profile construction. Cell types were annotated either manually or using the package “SingleR” (v1.6.1). For manual annotation, cells were clustered and assigned cell types based on the expression of cell type specific signature genes (SI.5a). Denoised results from InferCNV were used as the input (“infercnv.observations.txt”). The averaged copy number of each chromosome segment was calculated, and the difference between each cell’s copy number and the overall mean at each segment was calculated. The deviation was summed across the entire chromosome to obtain the distribution of the deviation. Cells sharing an absolute deviation that is more than 2.5 times standard deviation away from the distribution peak were marked as outliers for a certain chromosome in question.

### Transcriptomic gene sets

For analyzing both bulk and single-cell RNA-sequencing data sets, either hallmark gene sets (from the Human Molecular Signatures Database) or customized gene sets were used. Customized gene sets were used exclusively for macrophage-associated analyses and were made by combining gene sets from the following: Ahn et al., 2018; Cerezo-Wallis et al., 2020; Cunha et al., 2018; Mujal et al., 2022; Perry et al., 2019; Zhou et al., 2020.

### Statistical analysis & curve fitting

Statistical analyses and curve fitting were performed in GraphPad Prism 9.4. Details for each analysis are provided in the figure legends. Tumor and tumoroid growth curve data (projected area vs time) were fit to the exponential growth model (A = A_0_e^kt^ for tumors and A = A_1_e^k(t-1)^ for tumoroids) using non-linear least squares regression with prefactors A_0_ or A_1_ and *k*, the exponential growth rate. For differential gene expression analysis for both bulk RNA-sequencing and single-cell RNA-sequencing datasets, statistical analyses were done using either RStudio 2022.02.3+492 or Python 3.8.

## Code availability

Sequencing data were analyzed using RStudio 2022.02.3+492 and the Seurat package. Additional analyses after normalization were done on either RStudio or Python 3.8. No new code central to the conclusions of this study was developed.

## Author contributions

Conceptualization: BHH. Formal analysis: BHH, MW, HZ, SP, AC. Funding acquisition: BHH, JCA, LJD, DED. Investigation & data collection: BHH, MW, HZ, SP, LJD, AC, JCA, MPT, NMO, TM. Methodology: BHH, MW. Resources: DED. Visualization: BHH, MW. Writing: BHH, DED.

## Acknowledgements

This work was supported by funding from the following sources: NIH R01 HL124106 (DED), U54 CA193417 (DED), NSF GRFP DGE-1845298 (BHH, JCA, MPT), and NIH F32 CA228285 (LJD). The authors acknowledge the following University of Pennsylvania core facilities: Cell Center Stockroom, the Penn Cytomics and Cell Sorting Resource Laboratory, the Penn Genomic and Sequencing Core, and the Cell & Development Biology Microscopy Core.

## Competing interests

The authors declare no competing interests.

## Data availability statement

All data are available within the article and its supplementary information. Data can be provided upon reasonable request from the corresponding author.

---

**Supplementary Figure 1.**
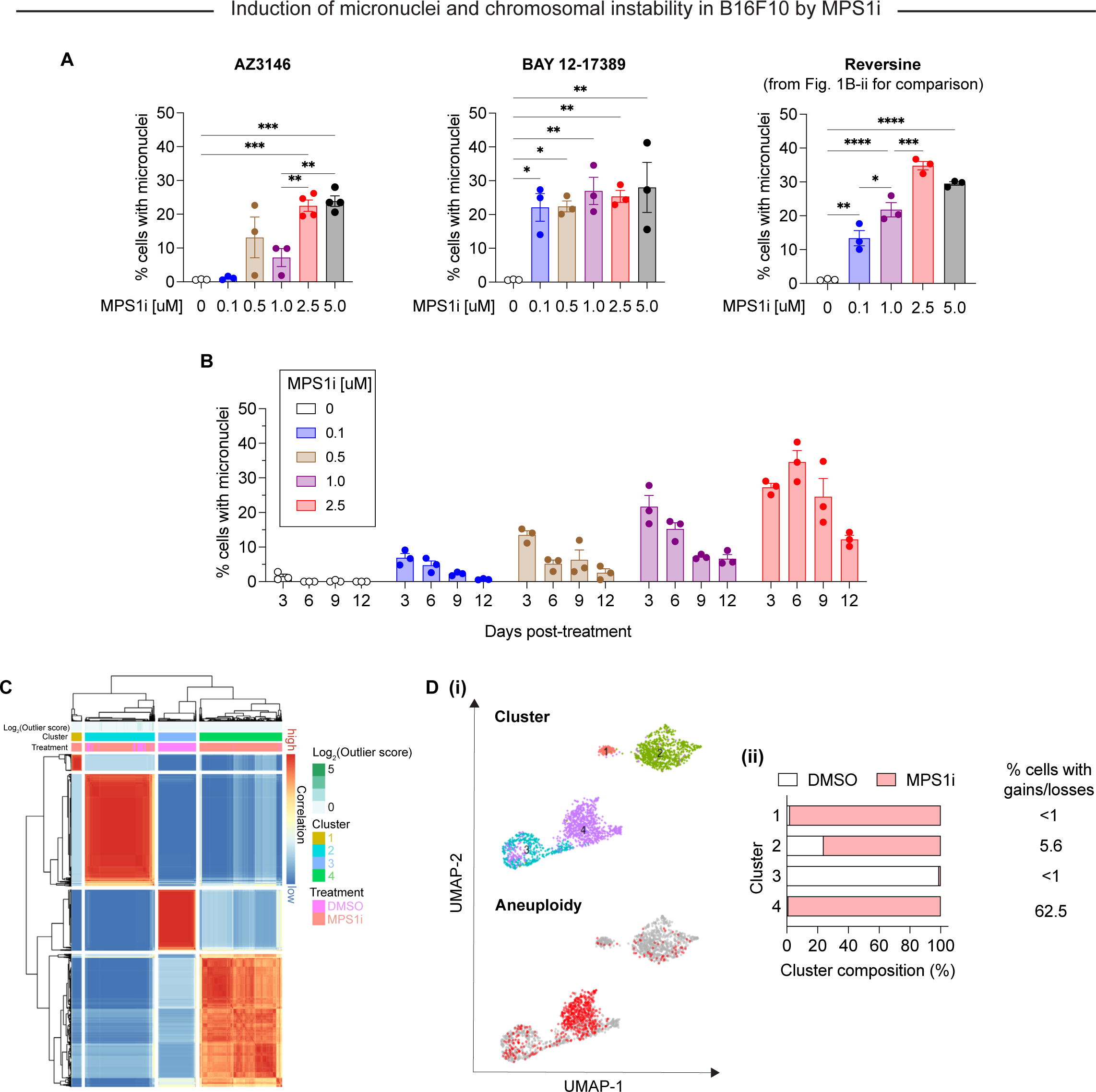
Characterization of MPS1i-induced genome and chromosomal instability in B16F10 mouse melanoma. **(A)** Quantification of micronuclei induced by different MPS1 inhibitors (AZ3146, BAY 12-17389, and reversine) at different concentrations. Statistical significance was calculated by ordinary one-way ANOVA and Tukey’s multiple comparison test (* p < 0.05; ** p < 0.01; *** p < 0.001; **** p < 0.0001). **(B)** Quantification of micronuclei induced by MPS1i up to 12 days after initial exposure, showing MPS1i yields micronuclei-positive persistent populations while untreated cells fail to produce micronuclei. **(C)** Consensus clustering of B16F10 analyzed in single-cell RNA-sequencing. B16F10 cells were treated with 2.5 μM MPS1i (reversine) or the equivalent volume of DMSO vehicle control. Cells were treated for 24 h, after which they were washed twice with PBS and allowed to recover for an additional 48 h. Cells were then collected and processes using 10X Genomics Chromium Single Cell Gene Expression kit for RNA isolation and library preparation. **(D) (i)** UMAP plots of expression profiles for all B16F10 analyzed in single-cell RNA-sequencing. Top: UMAP plots depict clusters in which B16F10 fall, as determined by consensus clustering. Bottom: UMAP plots highlight cells with detectable copy number variations (CNVs), as determined by InferCNV. Red circles indicate individual cells with at least one CNV. Gray circles represent individual cells with no detectable CNV, suggesting these are chromosomally stable. **(ii)** Quantification of the composition (MPS1i-treated or DMSO-treated B16F10) of each cluster.

**Supplementary Figure 2.**
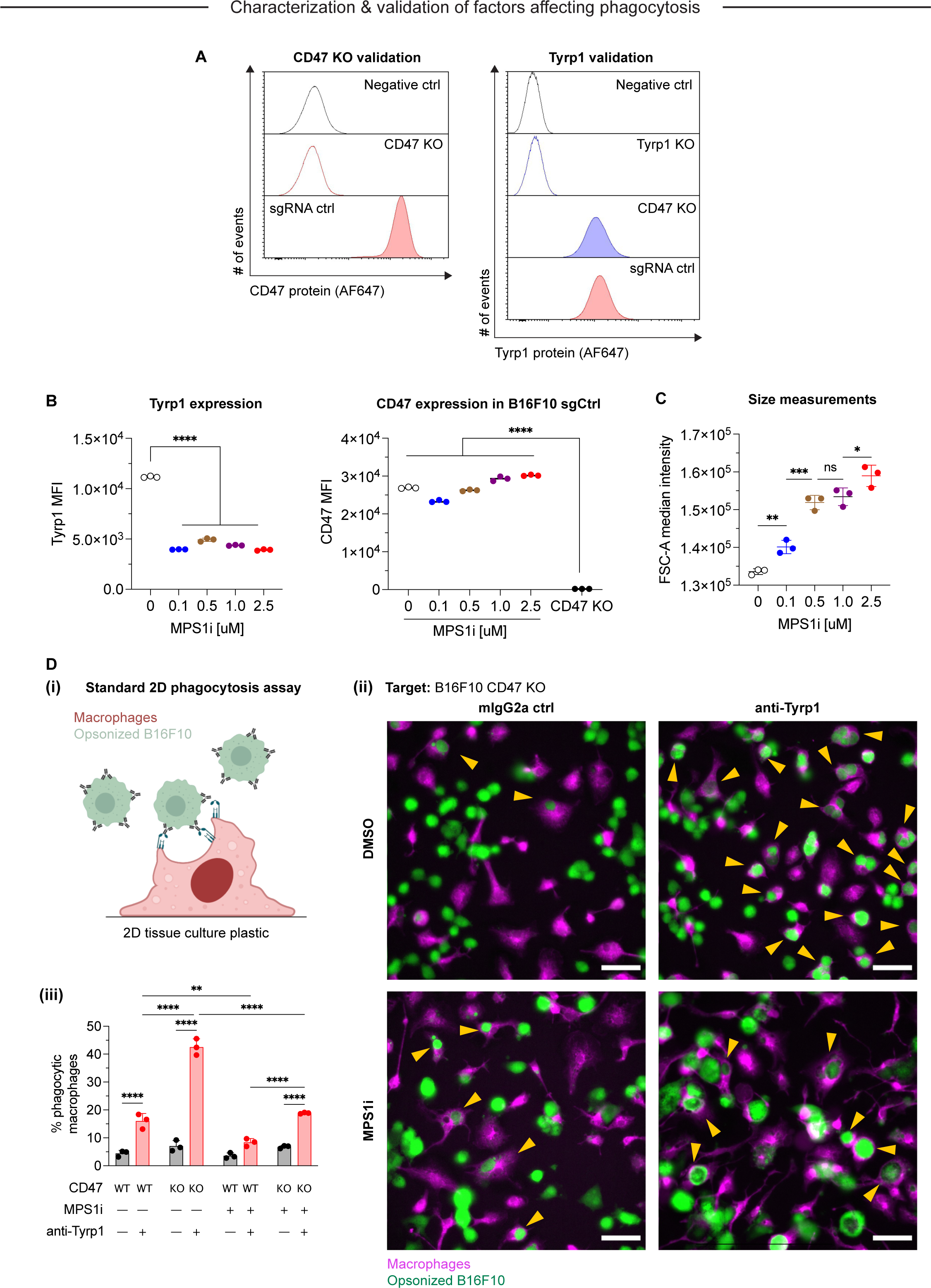
Characterization of MPS1i-induced genome and chromosomal instability in B16F10 mouse melanoma. **(A)** Flow cytometry analysis of CD47 and Tyrp1 expression on B16F10 mouse melanoma cells. Representative histograms for anti-CD47 (left) and anti-Tyrp1 (right) binding to B16F10 CD47 knockout (KO) and sgRNA ctrl (wild-type CD47 expression). Binding was detected by using secondary antibodies conjugated with Alexa Fluor 647. **(B)** Quantification of Tyrp1 expression by flow cytometry in both MPS1i- and DMSO-treated cells. Statistical significance was calculated by three-way ANOVA and Tukey’s multiple comparison test. Mean *±* SD shown, n = 3 replicates per condition (**** p < 0.0001). Quantification of CD47 expression by flow cytometry in both MPS1i- and DMSO-treated B16F10 sgCtrl cells. Statistical significance was calculated by three-way ANOVA and Tukey’s multiple comparison test. Mean *±* SD shown, n = 3 replicates per condition (**** p < 0.0001). **(C)** Quantification of cell size measurements by flow cytometry in both MPS1i- and DMSO-treated cells. Statistical significance was calculated by three-way ANOVA and Tukey’s multiple comparison test. Mean *±* SD shown, n = 3 replicates per condition (* p < 0.05; ** p < 0.01; *** p < 0.001). **(D)** Phagocytosis of B16F10 cells in in a standard phagocytosis assay on two-dimensional tissue culture plastic, with or without the following: IgG opsonization by anti-Tyrp1, CD47 KO, and MPS1i-treatment of B16F10. B16F10 cells were treated with MPS1i or DMSO prior to this assay, following the same protocol described in Fig. 1A. **(i)** Schematic of the 2D phagocytosis assay. **(ii)** Representative images of phagocytosis. B16F10 cells (green) were incubated with opsonizing anti-Tyrp1 or mouse IgG2a control and added to adherent bone marrow-derived macrophages (magenta). Random fields were imaged and then quantified to determine the percentage of phagocytic macrophages. Yellow arrowheads denote phagocytic events (complete engulfment of a target B16F10 cells). **(iii)** Quantification of phagocytic macrophages. Statistical significance was calculated by three-way ANOVA and Tukey’s multiple comparison test. Mean *±* SD shown, n = 3 replicates per condition (** p < 0.01; **** p < 0.0001).

**Supplementary Figure 3.**
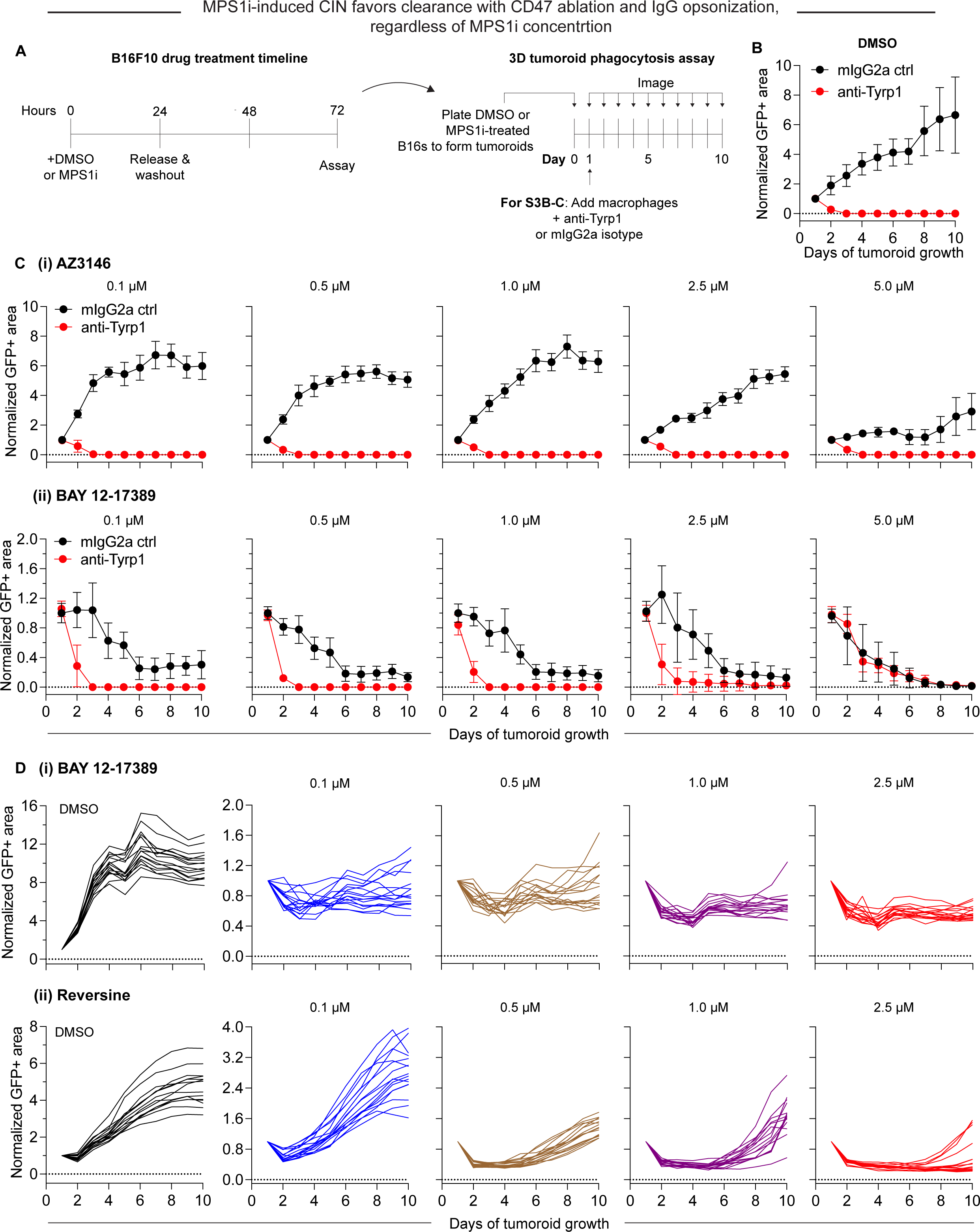
Macrophages readily clear CIN-afflicted tumoroids but only if CIN is accompanied by proliferation deficits.

**(A)** Timeline and schematic for generating engineered “immuno-tumoroids” for time-lapsed studies of macrophage-mediated phagocytosis of cancer cells. Tumoroids are formed by plating and culturing B16F10 cells on non-adhesive surfaces in U-bottom shaped wells. Prior to plating for tumoroid formation, B16F10 were treated with either MPS1i or DMSO (AZ3146, BAY 12-17389 or reversine). Tumoroid growth was measured by calculating the GFP+ area at the indicated timepoints (mean *±* SD, n = 16 total tumoroids from two independent experiments for each condition). All data were then normalized to average GFP+ area on day 1 of each drug treatment’s respective mouse IgG2a isotype control condition. Immuno-tumoroids are imaged at the listed timepoints. **(B)** To assess macrophage-mediated clearance, ∼24 h after plating, bone marrow-derived macrophages (BMDMs) with or without opsonizing anti-Tyrp1 are added to the cohesive B16F10 tumoroids. Tumoroid growth curves for DMSO-treated B16F10 CD47 knockout (KO) cells. **(C)** Tumoroid growth curves (with BMDMs) for MPS1i-treated B16F10 CD47 KO cells (treated with varying concentrations of MPS1i): **(i)** AZ3146, **(ii)** BAY 12-17389. **(D)** Tumoroid growth curves (without BMDMs) for MPS1i-treated B16F10 CD47 KO cells (treated with varying concentrations of MPS1i): **(i)** BAY 12-17389, **(ii)** reversine. Both BAY 12-17389 and reversine induce proliferation deficits, which may play an essential role in mediating clearance by macrophages that could not be observed in AZ3146-treated cells.

**Supplementary Figure 4.**
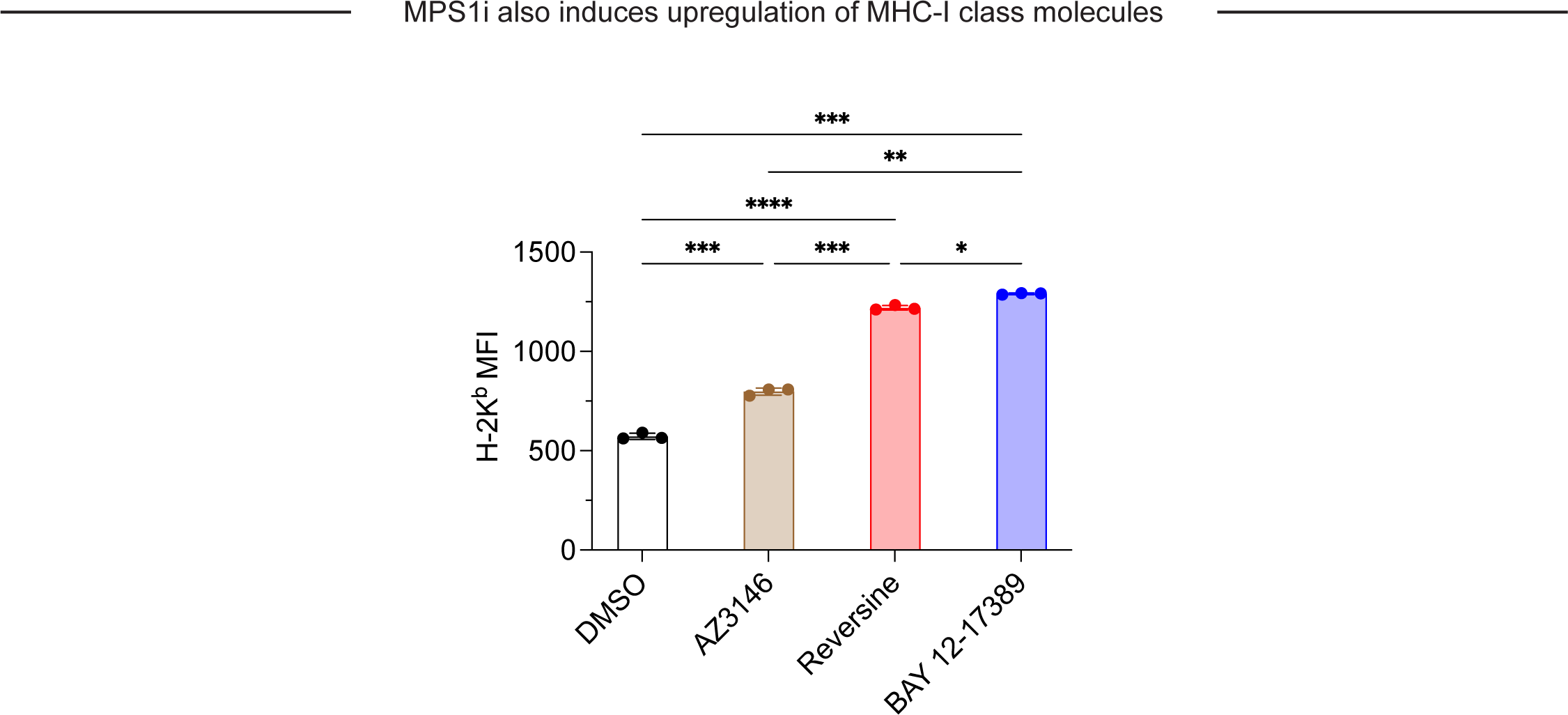
MPS1i-induced chromosomal instability upregulates MHC-1 class molecules on B16F10, suggesting increased antigen presentation. Flow cytometry analysis and quantification H-2K^b^ expression on B16F10 mouse melanoma cells. B16F10 cells were treated with MPS1i or the equivalent volume of DMSO vehicle control. Concentrations used: 2.5 μM AZ3146, 1.0 μM BAY 12-17380, and 2.5 μM reversine. Cells were treated for 24 h, after which they were washed twice with PBS and allowed to recover for an additional 48 h. Binding was detected by using primary antibodies conjugated with Alexa Fluor 647. Statistical significance was calculated by Brown-Forsythe and Welch ANOVA and Dunnett’s T3 multiple comparison test. Mean *±* SEM shown, n = 3 independent replicates (* p < 0.05; ** p < 0.01; *** p < 0.001; **** p < 0.0001).

**Supplementary Figure 5.**
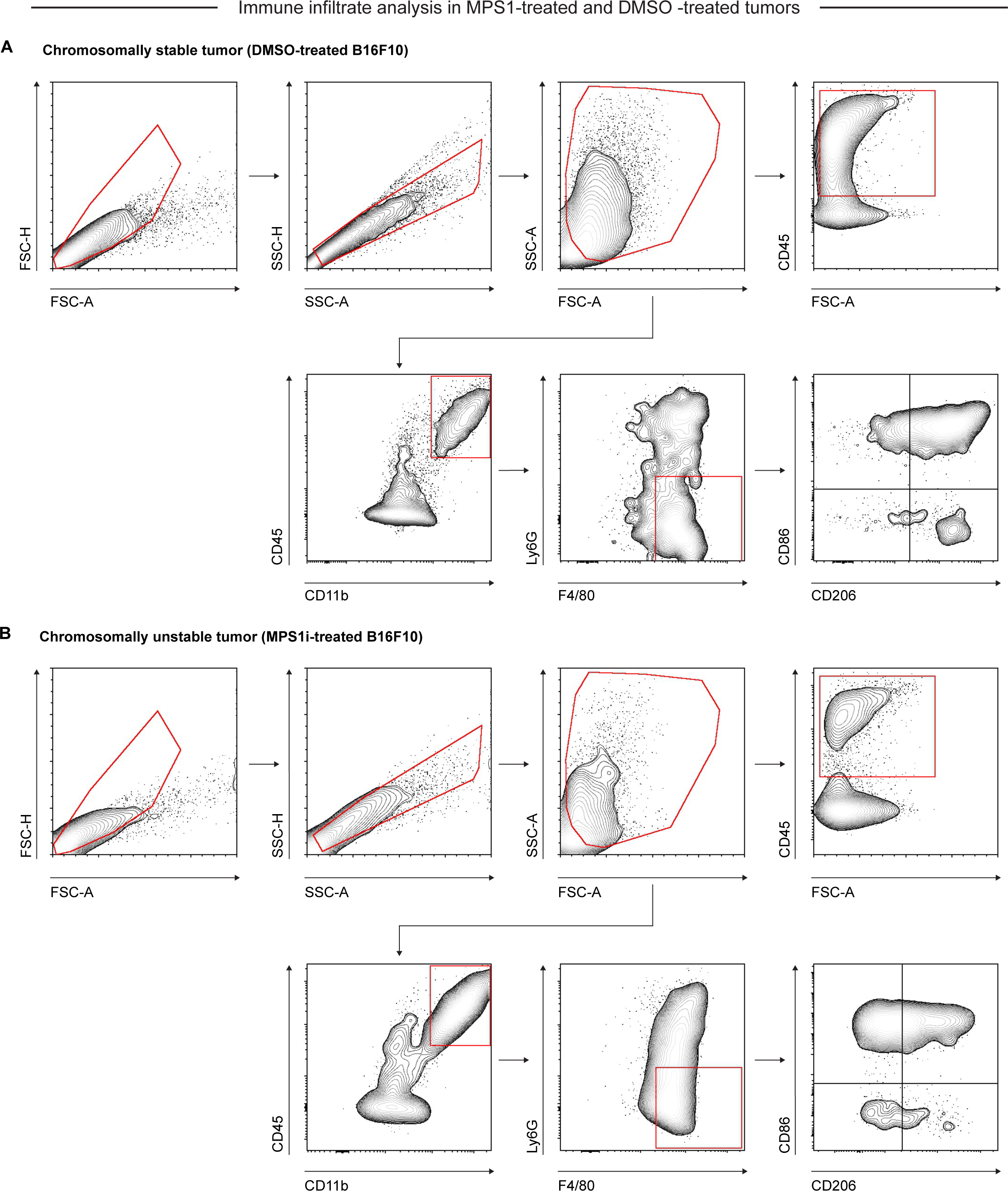
Flow cytometry gating strategy for identification & quantification of macrophage infiltrate and characterization in CIN-afflicted and chromosomally stable B16F10 tumors. Representative flow cytometry gating strategy for *in vivo* B16F10 CD47 KO tumor immune infiltrate five days after initial challenge. Tumors were comprised of cells treated with either **(i)** DMSO or **(ii)** MPS1i (2.5 μM), following the timeline schema in Fig. 2F. Singlets were separated from debris, doublets, and aggregates by FSC-A vs FSC-H and SSC-A vs SSC-H gates. Debris and dead cells were further removed by FSC-A vs SSC-A gating. Overall immune cell infiltrate was determined by CD45+ expression. Macrophages were then further identified based on Ly6G- and F4/80+ expression. Upon identification of macrophages, cells were further characterized for polarization based on CD86 (M1-like marker) and CD206 (M2-like marker) expression.

**Supplementary Figure 6.**
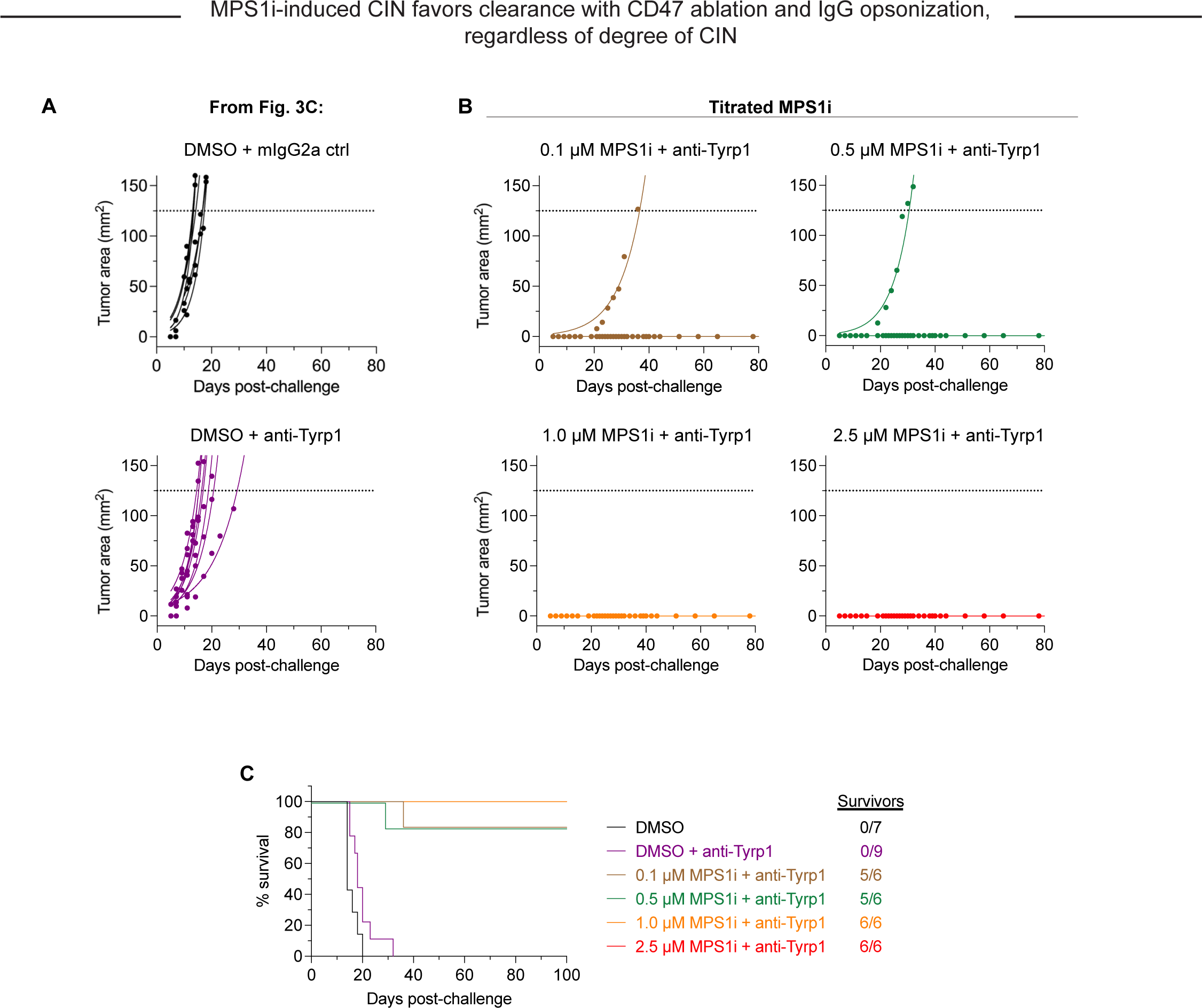
MPS1i-induced CIN favors clearance when paired with CD47 KO and IgG opsonization, regardless of the degree of CIN. Tumor growth curve of projected tumor area versus days after tumor challenge, with B16F10 CD47 KO cells. Each line represents a separate tumor and is fit with an exponential growth equation: A = A_0_e^kt^. Complete anti-tumor responses in which a tumor never grew are depicted with the same symbol as their growing counterparts and with solid lines at A = 0. **(A**) Tumor growth curve data in which there was no complete response, from Fig. 3C. n = 7 mice that were challenged with DMSO-treated B16F10 CD47 KO and subsequently treated with mouse IgG2a control, n = 9 mice that were challenged with DMSO-treated B16F10 CD47 KO and subsequently treated with anti-Tyrp1. **(B)** Tumor growth curve data from mice challenges with CIN-afflicted B16F10, but with varying concentration of MPS1i used to cause different degrees of instability. All mice here were challenged with B16F10 CD47 KO and treated with anti-Tyrp1 for conditions of maximal phagocytosis and to better compare to the ∼97% cure rate in Fig. 3C, 4D-ii. n = 6 mice challenged with cells with each tested MPS1i concentration: 0.1, 0.5, 1.0 or 2.5 μM. **(C)** Survival curves of mice up to 100 days after the tumor challenges in (A) and (B).

**Supplementary Figure 7.**
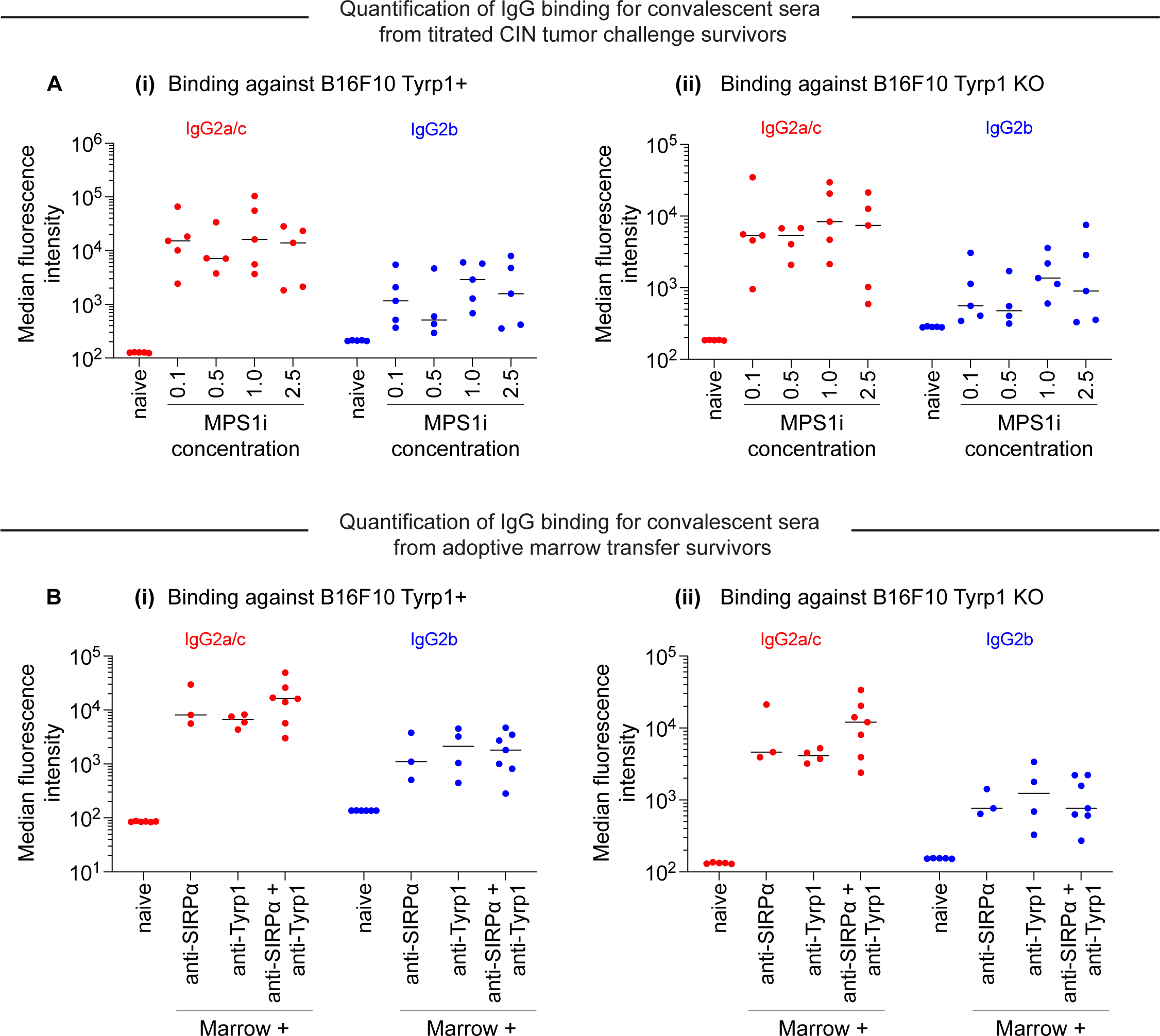
Survivors challenged with CIN-afflicted tumors generate anti-cancer IgG, regardless of the degree of CIN. Median fluorescence intensity quantification of IgG2a/c and IgG2b binding from sera from surviving mice with different degrees of chromosomal instability by titrating reversine concentration used to treat B16F10 CD47 KO cells before injection. Convalescent sera are collected from survivors from Fig. S6. Binding of IgG2a/c and IgG2b against **(A)** B16F10 cells expressing Tyrp1 and **(B)** B16F10 Tyrp1 knockout (KO). n = 4-5 distinct sera samples collected from survivors.

**Supplementary Figure 8.**
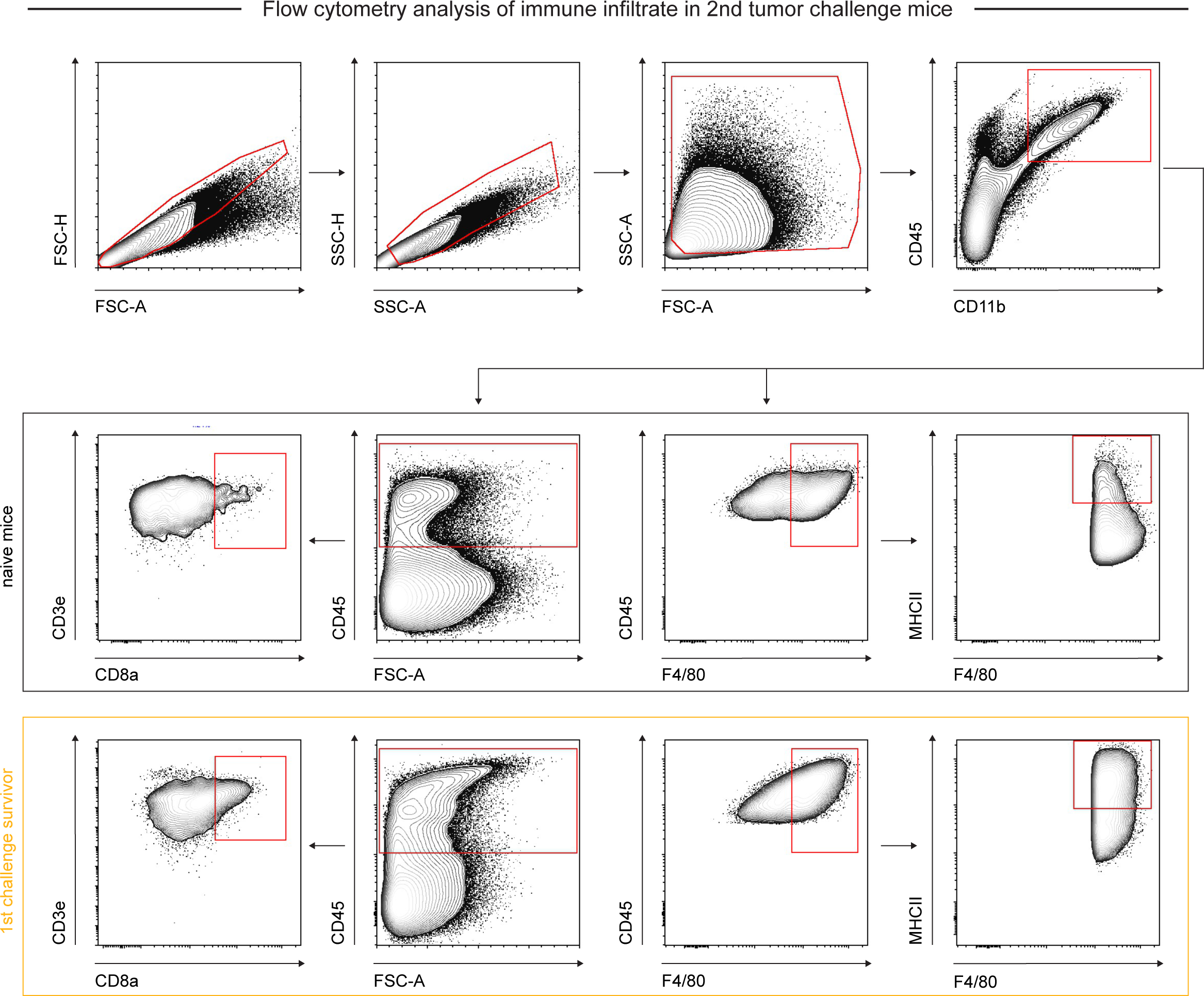
Flow cytometry gating strategy for identification & quantification of immune infiltrate and characterization in re-challenge experiments. Representative flow cytometry gating strategy for *in vivo* B16F10 CD47 KO tumor immune infiltrate from 2^nd^ challenge non-survivors and age-matched naïve controls. Singlets were separated from debris, doublets, and aggregates by FSC-A vs FSC-H and SSC-A vs SSC-H gates. Debris and dead cells were further removed by FSC-A vs SSC-A gating. For T cell quantification: CD45+ cells were gated on and then analyzed for CD3e and CD8a surface protein expression. For macrophage quantification: myeloid cells were isolated by CD45+ and CD11b+ expression, after which macrophages were isolated based on F4/80 expression.

**Supplementary Figure 9.**
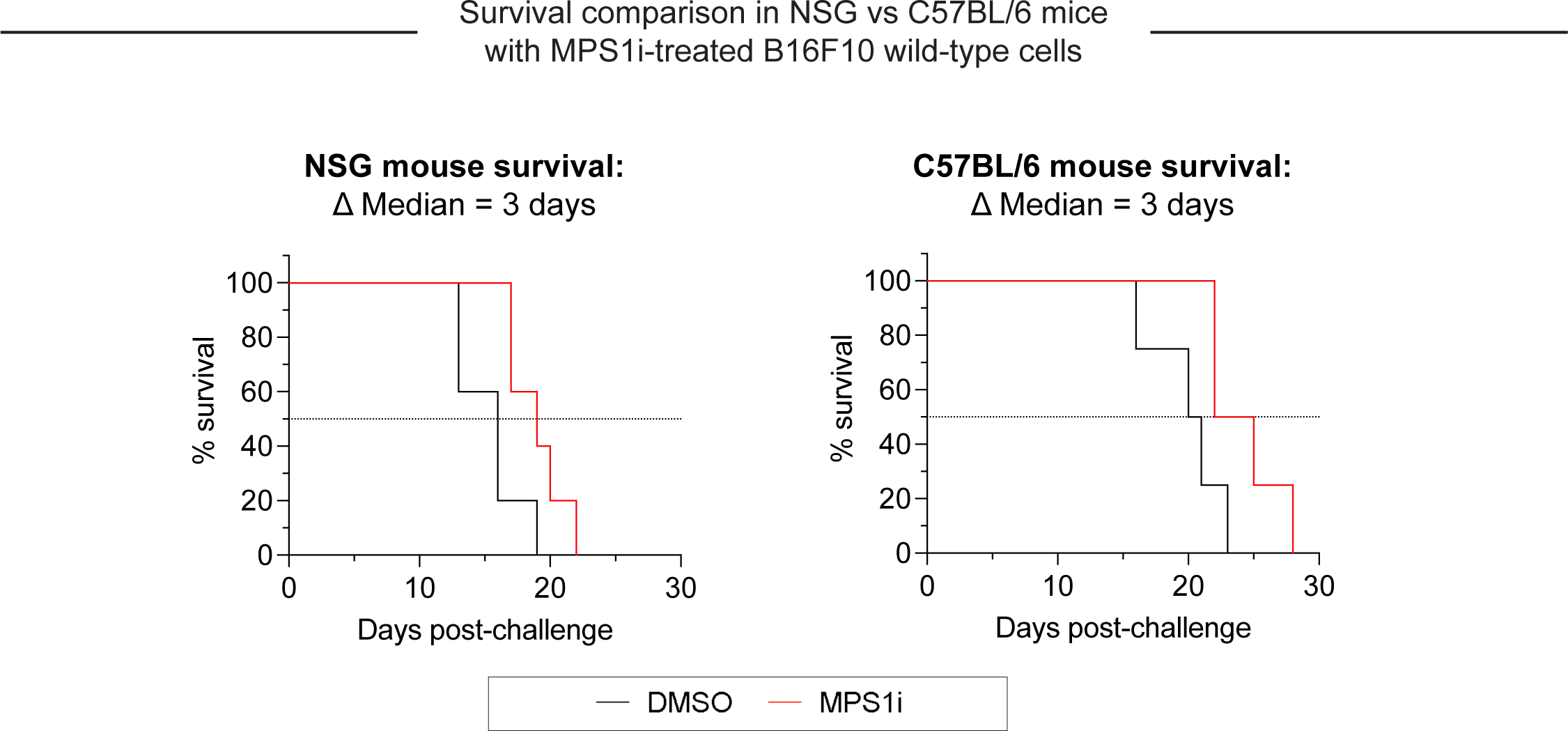
Growth of CIN-afflicted wild-type (WT) tumors in T- and B-cell deficient mice and T- and B-cell replete mice. Similar growth delays are found for MPS1i-pretreated B16F10 cells in T- and B-cell deficient NSG mice and immunocompetent C57BL/6 mice. Both types of mice have functional macrophages. Parallel studies *in vivo* were done with WT B16F10 ctrl cells cultured 24 h in 2.5 μM MPS1i (reversine or DMSO, then washed 3x in growth media for 5 min each and allowed to recover in growth media for 48 h. 200,000 cells in 100 uL PBS were injected subcutaneously into right flanks, and the standard size limit was used to determine survival curves. The C57BL/6 experiments were done independently here (by co-author L.J.D.) from the similar results (by B.H.H.) shown in Fig.3D-i, which provides evidence of reproducibility.

## Response for revised *eLife*

*Thank you for submitting your article “Chromosomal instability can favor macrophage-mediated immune response and induce a broad, vaccination-like anti-tumor IgG response” for consideration by eLife. Your article has been reviewed by 2 peer reviewers, and the assessment has been overseen by a Reviewing Editor and Carla Rothlin as the Senior Editor*.

### Response

We sincerely appreciate the time and efforts of the two Reviewers as well as the *eLife* Editors.

We apologize for the delay in submitting a revision, but multiple circumstances caused the delay. Most important was that a key mouse suite at Penn had bacterial/fungal contaminations which wiped out entire colonies, and shut down facilities for months. Although we eventually completed the requested experiments in the Fall, the first author of the manuscript wrote and defended his PhD thesis soon after the Reviewer comments were received, and his new position did not allow time until now to devote to the revision.

______

#### eLife assessment

*This study highlights a valuable finding that chromosomal instability can change immunes responses, in particular macrophages behaviours. The convincing results showing that the use of CD47 targeting and anti-Tyrp1 IgG can overcome changes in immune landscape in tumors and prolong survival of tumor-bearing mice. These findings reveal a new exciting dimension on how chromosomal instability can influence immune responses against tumor*.

#### Response

We thank the Editors for their enthusiasm and appreciation for this work. We also want to highlight our thanks for their careful reading, support, and patience while handling this manuscript. While this work provides useful insight into potential therapeutic implications of chromosomal instability in the macrophage immunotherapy field, we also hope it elucidates some novel basic science to further explore how chromosomal instability has such interesting effects on the immune system.

______

#### Public Reviews

#### Reviewer #1 (Public Review)

*The manuscript by Hayes et al. explored the potential of combining chromosomal instability with macrophage phagocytosis to enhance tumor clearance of B16-F10 melanoma. However, the manuscript suffers from substandard experimental design, some contradictory conclusions, and a lack of viable therapeutic effects*.

*The authors suggest that early-stage chromosomal instability (CIN) is a vulnerability for tumorigenesis, CD47-SIRPa interactions prevent effective phagocytosis, and opsonization combined with inhibition of the CD47-SIRPa axis can amplify tumor clearance. While these interactions are important, the experimental methodology used to address them is lacking*.

#### Reviewer #1 (Recommendations For The Authors)

*First, early stages of the tumor are essentially being defined as before implantation. In all cases, the tumor cells were pre-treated with MPS1i or had a genetic knockout of CD47. This makes it difficult to see how this would translate clinically*.

**Response:**

We greatly appreciate the Reviewer’s interest in the topic and its potential, but our manuscript makes no claims of immediate clinical translation. Chromosomal instability (CIN) studies have to date not yet discovered or described whether and how CIN can affect macrophage function. To our knowledge, this is the first study to begin such characterizations with various MPS1i drugs to induce CIN. Many variations of the approach can be envisioned for future studies.

Our Results include some key studies of cancer cells with wildtype levels of CD47-including in vivo tumor elimination (Fig.3E). Nonetheless, we do conduct some of our studies in a CD47 knockout context to remove this “brake” that generally impedes phagocytosis, with our goal being to better understand how CIN affects phagocytosis. As cited to some extent in our Introduction, there are many efforts in clinical trials to disrupt this macrophage checkpoint and others focused on macrophage immunotherapy. Whether CIN can be induced by clinically translatable drugs and specifically in cancer cells is beyond the scope of our studies.

*I would like to see the amount of CIN that occurs in WT B16F10 over the course of tumorigenesis (ie longer than 5 days). This is because I would assume that CIN would eventually occur in the WT B16-F10 regardless of whether MPS1i is being given. And if that’s the case, then the initiation of CIN at day 10 after implantation (for example) would still be considered “early stage” CIN. If the therapy is then initiated at this point, does the effect remain? Or put differently, how would the authors propose to induce the appropriate level of CIN in an established tumor? Why is pretreatment necessary?*

### Response

Untreated B16F10 cells fail to produce micronuclei over 12 days compared to MPS1i treated cells – as shown in a newly added panel in Fig. S1:

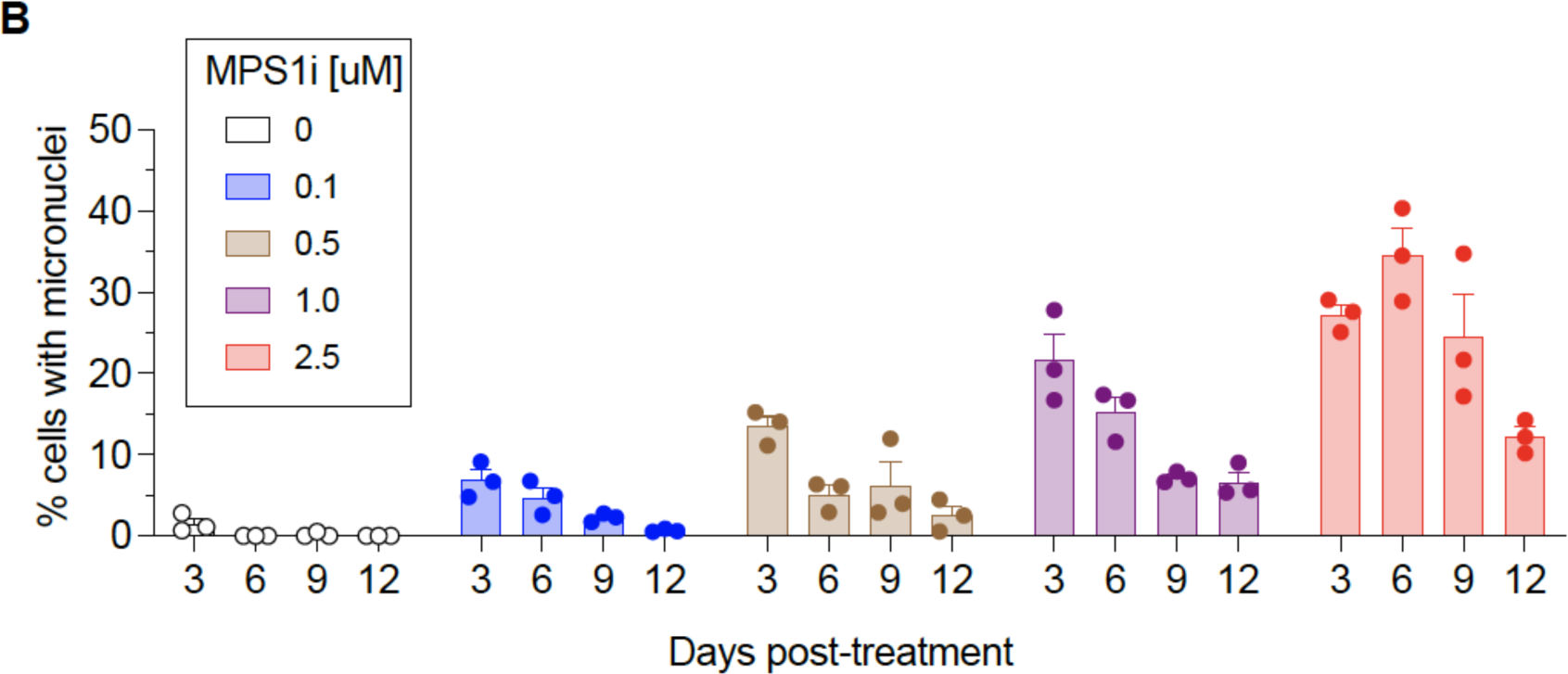

This helps support our decision to pre-treat cells with MPS1i to stimulate genomic instability and is described in the first section of Results:

“…we saw >10-fold increases of micronuclei over the cell line’s low basal level (∼1% of cells), and two other MPS1i inhibitors AZ3146 and BAY12-17389 confirm such effects (**Fig. S1A**). Micronuclei-positive cells can persist up to 12 days after treatment (**Fig. S1B**), while control cells maintain the low basal levels. The results suggest pre-treatment with MPS1i can simulate CIN in an experimental context even for 1-2 weeks, which may not typically occur at the same frequency during early tumor growth.

*It is known that PD-1 expression inhibits tumor-associated macrophage phagocytosis (Nature, 2017). Does MSP1i (sic) treatment affect the population of PD-1+ tumor macrophages in vivo?*

### Response

We thank the Reviewer for bringing up an interesting point.

Using the same tumor RNA-seq data that was used for Fig.1E, a heatmap of expression of PD-1 (gene *Pdcd1*) shows no consistent trend with MPS1i:

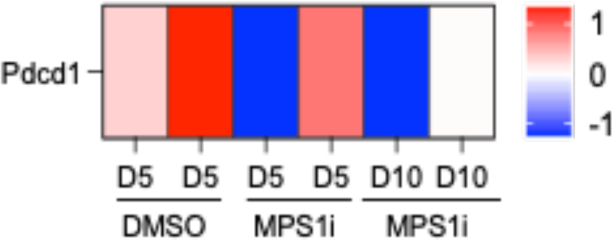

We also examined whether the secretome from CIN-afflicted cancer cells affect PD-1 expression in cultured macrophages, but we did not register any reads from our single-cell RNA-sequencing experiment for *Pdcd1* in any of the macrophage clusters from Fig. 1H.

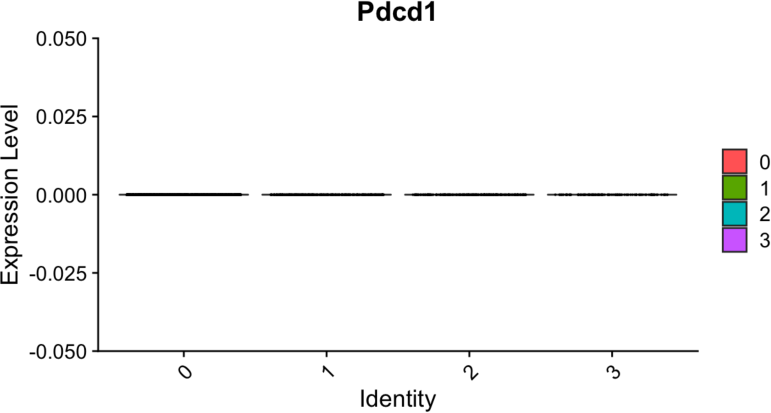

The Discussion section now includes a statement on this topic:

“…B16F10 tumors are poorly immunogenic, do not respond to either anti-CD47 or anti-PD-1/PD-L1 monotherapies, and show modest and variable cure rates (∼20-40%; Dooling et al., 2023; Hayes et al., 2023) even when macrophages have been made maximally phagocytic according to notions above. We should note here that our whole-tumor RNA-seq data (**Fig.1E**) shows expression of PD-1 (gene *Pdcd1*) follows no consistent trend upon MPS1i treatment, and that *Pdcd1* was not detected in our scRNA-seq data for macrophage cultures (**Fig.1G**) – motivating further study.”

*The authors must explain how the proposed therapy works since MPS1i increases tumor (cell) size, making it difficult for macrophages to phagocytose the tumor cells. It also reduces or suppresses Tyrp1 expression on the cancer cells, making it harder to opsonize. Since these were two main points for the rationale of this study, the authors need to reconcile them*.

### Response

We appreciate this comment and have re-organized this Results section to try to minimize confusion:

#### CIN-afflicted, CD47-knockout tumoroids are eliminated by Macrophages

To assess functional effects of macrophage polarization, we focused on a 3D “immuno-tumoroid” model in which macrophage activity can work (or not) over many days against a solid proliferating mass of cancer cells in non-adherent roundbottom wells (**Fig. 2A**) (Dooling et al., 2023). We used CD47 knockout (KO) B16F10 cells, which removes the inhibitory effect of CD47 on phagocytosis, noting that KO does not perturb surface levels of Tyrp1, which is targetable for opsonization with anti-Tyrp1 (**Fig. S2A**). BMDMs were added to pre-assembled tumoroids at a 3:1 ratio, and we first assessed surface protein expression of macrophage polarization markers. Consistent with our whole-tumor bulk RNA-sequencing and also single-cell RNA-sequencing of BMDM monocultures (**Fig. 1E, 1I-J**), BMDMs from immunotumoroids of MPS1i-treated B16F10 showed increased surface expression of M1-like markers MHCII and CD86 while showing decreased expression of M2-like markers CD163 and CD206 (**Fig. 2B-C**). Although these macrophages seemed poised for anticancer activity, the cancer cells showed decreased binding of anti-Tyrp1 (**Fig. S2B**) and ∼20% larger size in flow cytometry (**Fig. S2C**). The latter likely reflects cytokinesis defects and poly-ploidy as acute effects of CIN induction (Chunduri & Storchová, 2019; Mallin et al., 2022). Such cancer cell changes might explain why standard 2D phagocytosis assays show BMDMs attached to rigid plastic engulf relatively few anti-Tyrp1 opsonized cancer cells pretreated with MPS1i versus DMSO (**Fig. S2D**). In such cultures, BMDMs use their cytoskeleton to attach and spread, competing with engulfment of large and poorly opsonized targets. Noting that tumors in vivo are not as rigid as plastic, our 3D immunotumoroids eliminate attachment to plastic, and large numbers of macrophages can cluster and cooperate in engulfing cancer cells in a cohesive mass (Dooling et al., 2023). We indeed find CIN-afflicted tumoroids are eliminated by BMDMs regardless of anti-Tyrp1 opsonization (**Fig. 2D-E**), whereas anti-Tyrp1 is required for clearance of DMSO control tumoroids (**Fig. 2D, S3B**). Imaging also suggests that cancer CIN stimulates macrophages to cluster (compare Day-4 in **Fig. 2D**), which favors cooperative phagocytosis of tumoroids (Dooling et al., 2023), and occurs despite the lack of cancer cell opsonization and their larger cell size. The 3D immunotumoroid results with induced CIN are thus consistent with a more pro-phagocytic M1-type polarization (**Fig.1J and 2B,C**).

*The authors used varying numbers of tumor cells for the in vivo portions of the study; the first half of the manuscript uses 500,000 cells, while the latter half uses 200,000 cells. Why?*

### Response

The reasons for the difference in numbers is now clarified in the Methods:

For assessing immune infiltrates in early stages of tumor engraftment, when tumors are still small, we used a relatively high number of tumor cells (500,000 cells in Fig. 1D and Fig. 2F-G) to achieve sufficient cell numbers after dissociating the tumors, particularly for the slow-growing MPS1i-treated tumors. More specifically, with dissection, collagenase treatment, passage through a filter to remove clumps, we would lose many cells, and yet needed 100,000 viable cells or more for bulk RNA-seq suspensions and for flow cytometry measurements. For all other studies, 200,000 cancer cells were injected,

*The authors need to report the tumor volumes and the total number of cells isolated from the day five tumors to avoid grossly inflating the effect (i.e. Fig 2G and 4G)*.

### Response

We have added relevant numbers in the Methods:

For day 5 post-challenge measurements, 100,000 to 200,000 live cells were collected. For *in vivo* tumor infiltrate studies in re-challenged mice, 10 million live cells were collected.

Also, regarding tumor sizes and cell numbers, we have previously published relevant measurements in assessments of tumor growth. Please see:

Brandon H Hayes, Hui Zhu, Jason C Andrechak, Lawrence J Dooling, Dennis E Discher, Titrating CD47 by mismatch CRISPR-interference reveals incomplete repression can eliminate IgG-opsonized tumors but limits induction of antitumor IgG, *PNAS Nexus*, Volume 2, Issue 8, August 2023, pgad243, https://doi.org/10.1093/pnasnexus/pgad243

Dooling, L.J., Andrechak, J.C., Hayes, B.H. *et al*. Cooperative phagocytosis of solid tumours by macrophages triggers durable anti-tumour responses. *Nat. Biomed. Eng* 7, 1081–1096 (2023). https://doi.org/10.1038/s41551-023-01031-3

In the present study, similar tumor growth curves are provided for transparency, but the Kaplan-Meier curves as the key pieces of data in Fig. 3-4. Lastly, regarding reporting total cell number harvested, we based our experiments on previously accepted measurements that also reported numbers out of total harvested cells. See:

Cerezo-Wallis, D., Contreras-Alcalde, M., … Soengas, M.S., 2020. Midkine rewires the melanoma microenvironment toward a tolerogenic and immune-resistant state. Nat Med 26, 1865–1877. https://doi.org/10.1038/s41591-020-1073-3

*The figure titles need to be revised. For example, the title of Figure 1 claims that “MPS1i-induced chromosomal instability causes proliferation deficits in B16F10 tumors.” However, the evidence provided is weak. The authors only present GSEA analysis of proliferation and no functional evidence of impairment. The authors need to characterize this proliferation deficit using in vitro studies and functional studies of macrophage polarization. I would suggest proliferation assays (crystal violet, MTT, Incucyte, etc) to measure the B16 growth over time with MPS1i treatment*.

### Response

We thank the Reviewer for pointing this out. In Fig.1 we have minimized information regarding proliferation because it is later quantified in Figs.2D,E, S3, and 3D-i:

Fig.1F legend: Top downregulated hallmark gene sets in tumors comprised of MPS1i-treated B16F10 cells, showing downregulated DNA repair, cell cycle, and growth-related pathways, consistent with observations of slowed growth in culture and in vivo – as subsequently quantified.

*Then the authors could collect the tumor supernatant to culture with macrophages and determine polarization in vitro. I would also like to see functional studies of macrophage polarization (suppression assays, cytokine production, etc). Currently, the authors provide no functional studies*.

### Response

Fig.2B,C provides functional surface marker measurements of in vitro polarization toward anti-cancer M1 macrophages by MPS1i-pretreated tumor cells, consistent with gene expression in Fig.1G-J. Function is further shown as ant-cancer activity in Fig.2D,E, as now stated explicitly in the text:

“…In our 3D tumoroid *in vitro assays*, we found that macrophages can suppress the growth of chromosomally unstable tumoroids and clear them, surprisingly both with and without anti-Tyrp1 (**Fig. 2D-E**), regardless of MPS1i concentration used for treatment. Such a result is consistent with M1-type polarization (**Fig.1J and 2B,C**), which tends to be more pro-phagocytic. Such a result is consistent with M1-type polarization (**Fig.1J and 2B,C**), which tends to be more pro-phagocytic.”

*The authors claim that macrophages are the key effector cells, but they need to provide evidence for this claim*.

### Response

Other immune cells clearly contribute to the presented results because the IgG must eventually come from B cells. The text has been edited to indicate ‘macrophages are key <u>initiating</u>-effector cells’, and some evidence for this is the maximal survival of (WT B16 + Rev tumors) in Fig.3E upon treatment with Marrow Macrophages plus Macrophage-relevant SIRPa blockade and Macrophage-relevant IgG (via FcR). T cells do not have SIRPa or FcR.

*They can deplete macrophages and T and B cells to determine whether the effect remains or is ablated. This is the only definitive way to make this claim*.

### Response

To determine whether T and B cells might also be key <u>initiating</u>-effector cells, new experiments were done with mice depleted of T and B cells (per Fig.S9, below). We compared the growth of MPS1i vs DMSO treatments in these mice to results in mice with T and B cells (which should replicate our previous results in Fig.3D-i). We found that slower growth with Rev relative to DMSO was similar in mice without T and B cells compared to mice with T and B cells. We have added to the text our conclusion that: <u>T and B cells are *not* key initiating-effector cells</u>. Whereas B cells are effector cells at least in terms of eventually making anti-tumor IgG, our results show that macrophages are key <u>initiating</u>-effector cells because macrophages certainly affect the growth of (WT B16 + Rev tumors) when more are added (Fig.3E).

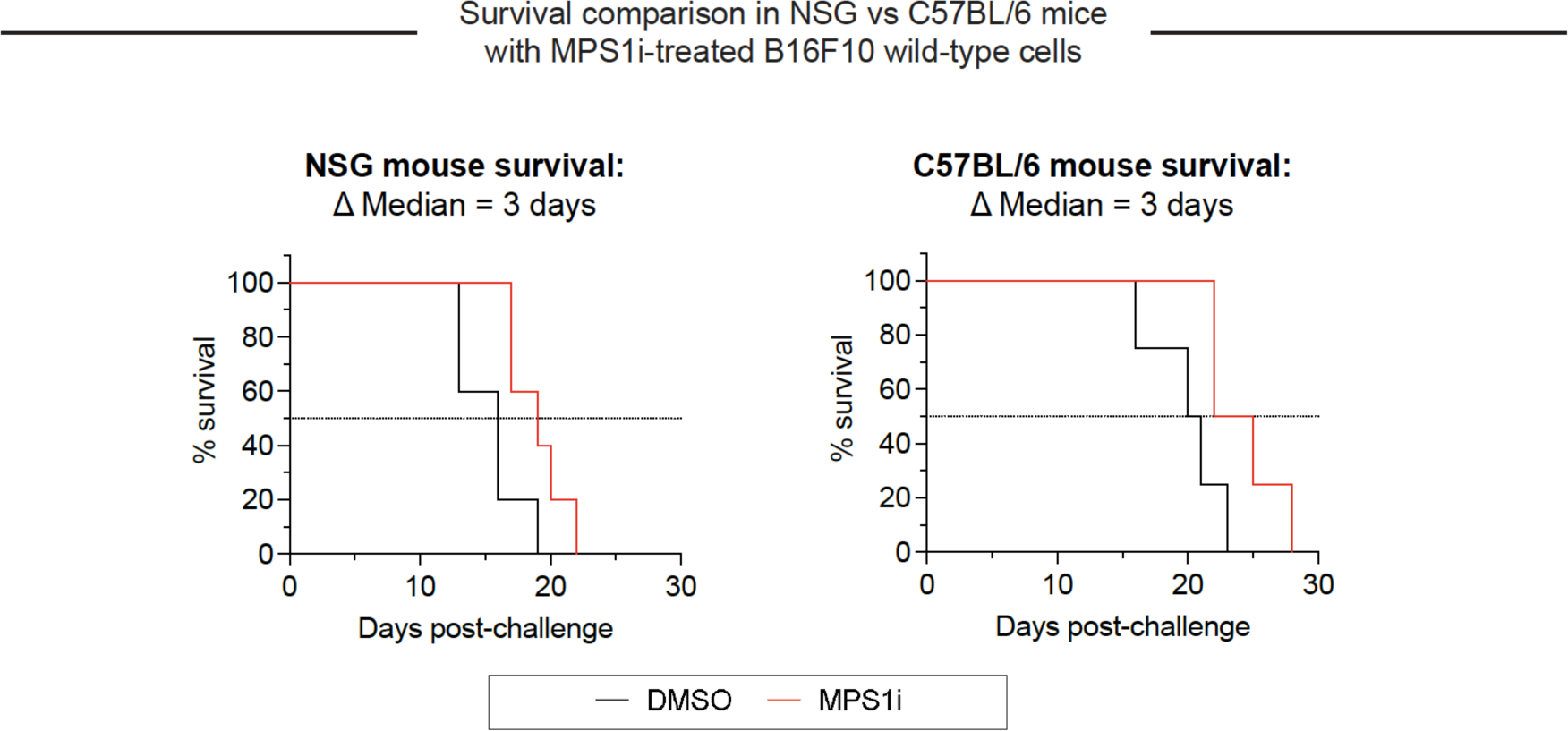

**Supplementary Figure 9. Growth of CIN-afflicted wild-type (WT) tumors in T- and B-cell deficient mice and T- and B-cell replete mice.**

Similar growth delays for MPS1i-pretreated B16F10 cells in T- and B-cell deficient NSG mice and immunocompetent C57BL/6 mice. Both types of mice have functional macrophages. Parallel studies *in vivo* were done with WT B16F10 ctrl cells cultured 24 h in 2.5 μM MPS1i (reversine or DMSO, then washed 3x in growth media for 5 min each and allowed to recover in growth media for 48 h. 200,000 cells in 100 uL PBS were injected subcutaneously into right flanks, and the standard size limit was used to determine survival curves. The C57BL/6 experiments were done independently here (by co-author L.J.D.) from the similar results (by B.H.H.) shown in Fig.3D-i, which provides evidence of reproducibility.

The Results section final paragraph describes all of this:

#### Reviewer #2 (Public Review)

*Harnessing macrophages to attack cancer is an immunotherapy strategy that has been steadily gaining interest. Whether macrophages alone can be powerful enough to permanently eliminate a tumor is a high-priority question. In addition, the factors making different tumors more vulnerable to macrophage attack have not been completely defined. In this paper, the authors find that chromosomal instability (CIN) in cancer cells improves the effect of macrophage targeted immunotherapies. They demonstrate that CIN tumors secrete factors that polarize macrophages to a more tumoricidal fate through several methods. The most compelling experiment is transferring conditioned media from MSP1 inhibited and control cancer cells, then using RNAseq to demonstrate that the MSP1-inhibited conditioned media causes a shift towards a more tumoricidal macrophage phenotype. In mice with MSP1 inhibited (CIN) B16 melanoma tumors, a combination of CD47 knockdown and anti-Tyrp1 IgG is sufficient for long term survival in nearly all mice. This combination is a striking improvement from conditions without CIN*.

*Like any interesting paper, this study leaves several unanswered questions. First, how do CIN tumors repolarize macrophages? The authors demonstrate that conditioned media is sufficient for this repolarization, implicating secreted factors, but the specific mechanism is unclear. In addition, the connection between the broad, vaccination-like IgG response and CIN is not completely delineated. The authors demonstrate that mice who successfully clear CIN tumors have a broad anti-tumor IgG response. This broad IgG response has previously been demonstrated for tumors that do not have CIN. It is not clear if CIN specifically enhances the anti-tumor IgG response or if the broad IgG response is similar to other tumors. Finally, CIN is always induced with MSP1 inhibition. To specifically attribute this phenotype to CIN it would be most compelling to demonstrate that tumors with CIN unrelated to MSP1 inhibition are also able to repolarize macrophages*.

*Overall, this is a thought-provoking study that will be of broad interest to many different fields including cancer biology, immunology and cell biology*.

### Response

We thank the Reviewer for their enthusiastic and positive comments toward the manuscript.

Our main purpose with this study has been discovery science oriented and mechanistic, with implications for improving macrophage immunotherapies. More experimentation needs to be done to further understand how this positive immune response emerges. However, we could address whether CIN enhances or not the anti-tumor IgG response by quantitative comparisons to our two other recent studies, and we conclude that it does not per new edits in the Abstract and the Results. See attached PPT for full details and comparison.

Abstract:

“CIN does not greatly affect the level of the induced response but does significantly increase survival.”

“…these results demonstrate induction of a generally potent anti-cancer antibody response to CIN-afflicted B16F10 in a CD47 KO context. Importantly, comparing these sera results for CIN-afflicted tumors to our recent studies of the same tumor model without CIN (Dooling et al., 2022; Hayes et al., 2022), we find similar levels of IgG induction (e.g. ∼100-fold above naive on average for IgG2a/c), similar increases in phagocytosis by sera opsonization (e.g. equivalent to anti-Tyrp1), and similar levels of suppressed tumoroid growth – including the variability.

…

However, median survival increased (21 days) compared to their naïve counterparts (14 days), supporting the initial hypothesis of prolonged survival and consistent not only with past results indicating major benefits of a prime-&-boost approach with anti-Tyrp1 (Dooling et al., 2022) but also with the noted similarities in induced IgG levels.”

Future studies could certainly focus on trying to identify what secreted factors might be inducing the M1-like polarization (using ELISA assays for cytokine detection, for example). This could be important because a main finding here is that we achieve nearly a 100% success rate in clearing tumors when we combine CD47 ablation and IgG opsonization with cancer cell CIN. Previous studies were only able to achieve about 40% cures in mice when working with CD47 disription and IgG opsonization alone, suggesting CIN in this experimental context does improve macrophage response.

Lastly, we agree with the Reviewer that future studies should also address how CIN in general (not MPS1i-induced) affects tumor growth. The final paragraph of our Discussion at least cites support for consistent effects of M1-like polarization:

“The effects of CIN and aneuploidy in macrophages certainly requires further investigation. We did publish recently that M1-like polarization of BMDMs with IFNγ priming is sufficient to suppress growth of B16 tumoroids with anti-Tyrp1 opsonization more rapidly than unpolarized/unprimed macrophages and much more rapidly than M2-like polarization of BMDMs with IL4 (Extended Data Fig.5a in Dooling et al., 2023); hence, anti-cancer polarization contributes in this assay. While the secretome from MPS1i-treated cancer cells has been found to trigger…”

Nonetheless, we can only speculate that there is a threshold of CIN reached by a certain timepoint in tumor engraftment and growth. Natural CIN might not be enough, so we pursued a pharmacological approach consistent with ongoing pre-clinical studies (https://doi.org/10.1158/1535-7163.MCT-15-0500). Future studies should consider trying knockdown models to gradually accrue CIN in tumors or using more relevant pharmacological drugs that are known to induce CIN not associated with the spindle. We believe, however, that these are larger questions on their own and are beyond the scope of the foundational discoveries in this manuscript.

#### Reviewer #2 (Recommendations For The Authors)

*None*

### Reponse

We again thank the Reviewer for their support and enthusiasm for the manuscript. We made some additional changes and more data to address questions posed by the other Reviewer that we hope you find to help the manuscript further.

## Notes

### Competing Interest Statement

The authors have declared no competing interest.

### Summary of Updates

Edits address eLife Reviewer comments and include new experimental results and analyses.

## References

Ahn, J., Xia, T., Rabasa Capote, A., Betancourt, D., Barber, G.N., 2018. Extrinsic Phagocyte-Dependent STING Signaling Dictates the Immunogenicity of Dying Cells. Cancer Cell 33, 862–873.e5. 10.1016/j.ccell.2018.03.027

Alvey, C.M., Spinler, K.R., Irianto, J., Pfeifer, C.R., Hayes, B., Xia, Y., Cho, S., Dingal, P.C.P.D., Hsu, J., Smith, L., Tewari, M., Discher, D.E., 2017. SIRPA-Inhibited, Marrow-Derived Macrophages Engorge, Accumulate, and Differentiate in Antibody-Targeted Regression of Solid Tumors. Current Biology 27, 2065–2077.e6. 10.1016/j.cub.2017.06.005

Dooling, L.J., Andrechak, J.C., Hayes, B.H., Kadu, S., Zhang, W., Pan, R., Vashisth, M., Irianto, J., Alvey, C.M., Ma, L., Discher, D.E., 2023. Cooperative phagocytosis of solid tumours by macrophages triggers durable anti-tumour responses. Nat. Biomed. Eng 7, 1081–1096. 10.1038/s41551-023-01031-3

Andrechak, J.C., Dooling, L.J., Tobin, M.P., Zhang, W., Hayes, B.H., Lee, J.Y., Jin, X., Irianto, J., Discher, D.E., 2022. CD47-SIRPα Checkpoint Disruption in Metastases Requires Tumor-Targeting Antibody for Molecular and Engineered Macrophage Therapies. Cancers 14, 1930. 10.3390/cancers14081930

Ben-David, U., Amon, A., 2020. Context is everything: aneuploidy in cancer. Nat Rev Genet 21, 44–62. 10.1038/s41576-019-0171-x

Boilève, A., Senovilla, L., Vitale, I., Lissa, D., Martins, I., Métivier, D., van den Brink, S., Clevers, H., Galluzzi, L., Castedo, M., Kroemer, G., 2013. Immunosurveillance against tetraploidization-induced colon tumorigenesis. Cell Cycle 12, 473–479. 10.4161/cc.23369

Bruhns, P., 2012. Properties of mouse and human IgG receptors and their contribution to disease models. Blood 119, 5640–5649. 10.1182/blood-2012-01-380121

Cerezo-Wallis, D., Contreras-Alcalde, M., Troulé, K., Catena, X., Mucientes, C., Calvo, T.G., Cañón, E., Tejedo, C., Pennacchi, P.C., Hogan, S., Kölblinger, P., Tejero, H., Chen, A.X., Ibarz, N., Graña-Castro, O., Martinez, L., Muñoz, J., Ortiz-Romero, P., Rodriguez-Peralto, J.L., Gómez-López, G., Al-Shahrour, F., Rabadán, R., Levesque, M.P., Olmeda, D., Soengas, M.S., 2020. Midkine rewires the melanoma microenvironment toward a tolerogenic and immune-resistant state. Nat Med 26, 1865–1877. 10.1038/s41591-020-1073-3

Champion, J.A., Mitragotri, S., 2006. Role of target geometry in phagocytosis. Proc. Natl. Acad. Sci. U.S.A. 103, 4930–4934. 10.1073/pnas.0600997103

Chao, M.P., Jaiswal, S., Weissman-Tsukamoto, R., Alizadeh, A.A., Gentles, A.J., Volkmer, J., Weiskopf, K., Willingham, S.B., Raveh, T., Park, C.Y., Majeti, R., Weissman, I.L., 2010. Calreticulin Is the Dominant Pro-Phagocytic Signal on Multiple Human Cancers and Is Counterbalanced by CD47. Sci. Transl. Med. 2. 10.1126/scitranslmed.3001375

Chunduri, N.K., Storchová, Z., 2019. The diverse consequences of aneuploidy. Nat Cell Biol 21, 54–62. 10.1038/s41556-018-0243-8

Cohen-Sharir, Y., McFarland, J.M., Abdusamad, M., Marquis, C., Bernhard, S.V., Kazachkova, M., Tang, H., Ippolito, M.R., Laue, K., Zerbib, J., Malaby, H.L.H., Jones, A., Stautmeister, L.-M., Bockaj, I., Wardenaar, R., Lyons, N., Nagaraja, A., Bass, A.J., Spierings, D.C.J., Foijer, F., Beroukhim, R., Santaguida, S., Golub, T.R., Stumpff, J., Storchová, Z., Ben-David, U., 2021. Aneuploidy renders cancer cells vulnerable to mitotic checkpoint inhibition. Nature 590, 486–491. 10.1038/s41586-020-03114-6

Crasta, K., Ganem, N.J., Dagher, R., Lantermann, A.B., Ivanova, E.V., Pan, Y., Nezi, L., Protopopov, A., Chowdhury, D., Pellman, D., 2012. DNA breaks and chromosome pulverization from errors in mitosis. Nature 482, 53–58. 10.1038/nature10802

Cunha, L.D., Yang, M., Carter, R., Guy, C., Harris, L., Crawford, J.C., Quarato, G., Boada-Romero, E., Kalkavan, H., Johnson, M.D.L., Natarajan, S., Turnis, M.E., Finkelstein, D., Opferman, J.T., Gawad, C., Green, D.R., 2018. LC3-Associated Phagocytosis in Myeloid Cells Promotes Tumor Immune Tolerance. Cell 175, 429–441.e16. 10.1016/j.cell.2018.08.061

Davoli, T., Uno, H., Wooten, E.C., Elledge, S.J., 2017. Tumor aneuploidy correlates with markers of immune evasion and with reduced response to immunotherapy. Science 355, eaaf8399. 10.1126/science.aaf8399

Fernando, M.R., Reyes, J.L., Iannuzzi, J., Leung, G., McKay, D.M., 2014. The Pro-Inflammatory Cytokine, Interleukin-6, Enhances the Polarization of Alternatively Activated Macrophages. PLoS ONE 9, e94188. 10.1371/journal.pone.0094188

Georgouli, M., Herraiz, C., Crosas-Molist, E., Fanshawe, B., Maiques, O., Perdrix, A., Pandya, P., Rodriguez-Hernandez, I., Ilieva, K.M., Cantelli, G., Karagiannis, P., Mele, S., Lam, H., Josephs, D.H., Matias-Guiu, X., Marti, R.M., Nestle, F.O., Orgaz, J.L., Malanchi, I., Fruhwirth, G.O., Karagiannis, S.N., Sanz-Moreno, V., 2019. Regional Activation of Myosin II in Cancer Cells Drives Tumor Progression via a Secretory Cross-Talk with the Immune Microenvironment. Cell 176, 757–774.e23. 10.1016/j.cell.2018.12.038

Harding, S.M., Benci, J.L., Irianto, J., Discher, D.E., Minn, A.J., Greenberg, R.A., 2017. Mitotic progression following DNA damage enables pattern recognition within micronuclei. Nature 548, 466–470. 10.1038/nature23470

Hayes, B.H., Tsai, R.K., Dooling, L.J., Kadu, S., Lee, J.Y., Pantano, D., Rodriguez, P.L., Subramanian, S., Shin, J.-W., Discher, D.E., 2020. Macrophages show higher levels of engulfment after disruption of *cis* interactions between CD47 and the checkpoint receptor SIRPα. Journal of Cell Science 133, jcs237800. 10.1242/jcs.237800

Hayes, B.H., Zhu, H., Andrechak, J.C., Dooling, L.J., Discher, D.E., 2023. Titrating CD47 by mismatch CRISPR-interference reveals incomplete repression can eliminate IgG-opsonized tumors but limits induction of antitumor IgG. PNAS Nexus 2, pgad243. 10.1093/pnasnexus/pgad243

Ingram, J.R., Blomberg, O.S., Sockolosky, J.T., Ali, L., Schmidt, F.I., Pishesha, N., Espinosa, C., Dougan, S.K., Garcia, K.C., Ploegh, H.L., Dougan, M., 2017. Localized CD47 blockade enhances immunotherapy for murine melanoma. Proc. Natl. Acad. Sci. U.S.A. 114, 10184– 10189. 10.1073/pnas.1710776114

Jablonski, K.A., Amici, S.A., Webb, L.M., Ruiz-Rosado, J. de D., Popovich, P.G., Partida-Sanchez, S., Guerau-de-Arellano, M., 2015. Novel Markers to Delineate Murine M1 and M2 Macrophages. PLoS ONE 10, e0145342. 10.1371/journal.pone.0145342

Jalil, A.R., Andrechak, J.C., Discher, D.E., 2020. Macrophage checkpoint blockade: results from initial clinical trials, binding analyses, and CD47-SIRPα structure–function. Antibody Therapeutics 3, 80–94. 10.1093/abt/tbaa006

Kamber, R.A., Nishiga, Y., Morton, B., Banuelos, A.M., Barkal, A.A., Vences-Catalán, F., Gu, M., Fernandez, D., Seoane, J.A., Yao, D., Liu, K., Lin, S., Spees, K., Curtis, C., Jerby-Arnon, L., Weissman, I.L., Sage, J., Bassik, M.C., 2021. Inter-cellular CRISPR screens reveal regulators of cancer cell phagocytosis. Nature 597, 549–554. 10.1038/s41586-021-03879-4

Kitajima, S., Tani, T., Springer, B.F., Campisi, M., Osaki, T., Haratani, K., Chen, M., Knelson, E.H., Mahadevan, N.R., Ritter, J., Yoshida, R., Köhler, J., Ogino, A., Nozawa, R.-S., Sundararaman, S.K., Thai, T.C., Homme, M., Piel, B., Kivlehan, S., Obua, B.N., Purcell, C., Yajima, M., Barbie, T.U., Lizotte, P.H., Jänne, P.A., Paweletz, C.P., Gokhale, P.C., Barbie, D.A., 2022. MPS1 inhibition primes immunogenicity of KRAS-LKB1 mutant lung cancer. Cancer Cell 40, 1128–1144.e8. 10.1016/j.ccell.2022.08.015

Krysko, D.V., Ravichandran, K.S., Vandenabeele, P., 2018. Macrophages regulate the clearance of living cells by calreticulin. Nat Commun 9, 4644. 10.1038/s41467-018-06807-9

Mackenzie, K.J., Carroll, P., Martin, C.-A., Murina, O., Fluteau, A., Simpson, D.J., Olova, N., Sutcliffe, H., Rainger, J.K., Leitch, A., Osborn, R.T., Wheeler, A.P., Nowotny, M., Gilbert, N., Chandra, T., Reijns, M.A.M., Jackson, A.P., 2017. cGAS surveillance of micronuclei links genome instability to innate immunity. Nature 548, 461–465. 10.1038/nature23449

Mallin, M.M., Kim, N., Choudhury, M.I., Lee, S.J., An, S.S., Sun, S.X., Konstantopoulos, K., Pienta, K.J., Amend, S.R., 2022. Cells in the Polyaneuploid Cancer Cell (PACC) state have increased metastatic potential (preprint). Cancer Biology. 10.1101/2022.09.16.508155

Morrissey, M.A., Kern, N., Vale, R.D., 2020. CD47 Ligation Repositions the Inhibitory Receptor SIRPA to Suppress Integrin Activation and Phagocytosis. Immunity 53, 290–302.e6. 10.1016/j.immuni.2020.07.008

Mujal, A.M., Combes, A.J., Rao, A.A., Binnewies, M., Samad, B., Tsui, J., Boissonnas, A., Pollack, J.L., Argüello, R.J., Meng, M.V., Porten, S.P., Ruhland, M.K., Barry, K.C., Chan, V., Krummel, M.F., 2022. Holistic Characterization of Tumor Monocyte-to-Macrophage Differentiation Integrates Distinct Immune Phenotypes in Kidney Cancer. Cancer Immunology Research 10, 403–419. 10.1158/2326-6066.CIR-21-0588

Nia, H.T., Munn, L.L., Jain, R.K., 2020. Physical traits of cancer. Science 370, eaaz0868. 10.1126/science.aaz0868

Nimmerjahn, F., Lux, A., Albert, H., Woigk, M., Lehmann, C., Dudziak, D., Smith, P., Ravetch, J.V., 2010. FcγRIV deletion reveals its central role for IgG2a and IgG2b activity in vivo. Proc. Natl. Acad. Sci. U.S.A. 107, 19396–19401. 10.1073/pnas.1014515107

Noy, R., Pollard, J.W., 2014. Tumor-Associated Macrophages: From Mechanisms to Therapy. Immunity 41, 49–61. 10.1016/j.immuni.2014.06.010

Oldenborg, P.-A., Zheleznyak, A., Fang, Y.-F., Lagenaur, C.F., Gresham, H.D., Lindberg, F.P., 2000. Role of CD47 as a Marker of Self on Red Blood Cells. Science 288, 2051–2054. 10.1126/science.288.5473.2051

Perry, J.S.A., Morioka, S., Medina, C.B., Iker Etchegaray, J., Barron, B., Raymond, M.H., Lucas, C.D., Onengut-Gumuscu, S., Delpire, E., Ravichandran, K.S., 2019. Interpreting an apoptotic corpse as anti-inflammatory involves a chloride sensing pathway. Nat Cell Biol 21, 1532–1543. 10.1038/s41556-019-0431-1

Santaguida, S., Richardson, A., Iyer, D.R., M’Saad, O., Zasadil, L., Knouse, K.A., Wong, Y.L., Rhind, N., Desai, A., Amon, A., 2017. Chromosome Mis-segregation Generates Cell-Cycle-Arrested Cells with Complex Karyotypes that Are Eliminated by the Immune System. Developmental Cell 41, 638–651.e5. 10.1016/j.devcel.2017.05.022

Senovilla, L., Vitale, I., Martins, I., Tailler, M., Pailleret, C., Michaud, M., Galluzzi, L., Adjemian, S., Kepp, O., Niso-Santano, M., Shen, S., Mariño, G., Criollo, A., Boilève, A., Job, B., Ladoire, S., Ghiringhelli, F., Sistigu, A., Yamazaki, T., Rello-Varona, S., Locher, C., Poirier-Colame, V., Talbot, M., Valent, A., Berardinelli, F., Antoccia, A., Ciccosanti, F., Fimia, G.M., Piacentini, M., Fueyo, A., Messina, N.L., Li, M., Chan, C.J., Sigl, V., Pourcher, G., Ruckenstuhl, C., Carmona-Gutierrez, D., Lazar, V., Penninger, J.M., Madeo, F., López-Otín, C., Smyth, M.J., Zitvogel, L., Castedo, M., Kroemer, G., 2012. An Immunosurveillance Mechanism Controls Cancer Cell Ploidy. Science 337, 1678–1684. 10.1126/science.1224922

Sockolosky, J.T., Dougan, M., Ingram, J.R., Ho, C.C.M., Kauke, M.J., Almo, S.C., Ploegh, H.L., Garcia, K.C., 2016. Durable antitumor responses to CD47 blockade require adaptive immune stimulation. Proc. Natl. Acad. Sci. U.S.A. 113. 10.1073/pnas.1604268113

Spurr, L.F., Martinez, C.A., Kang, W., Chen, M., Zha, Y., Hseu, R., Gutiontov, S.I., Turchan, W.T., Lynch, C.M., Pointer, K.B., Chang, P., Murgu, S., Husain, A.N., Cody, B., Vokes, E.E., Bestvina, C.M., Patel, J.D., Diehn, M., Gajewski, T.F., Weichselbaum, R.R., Chmura, S.J., Pitroda, S.P., 2022a. Highly aneuploid non-small cell lung cancer shows enhanced responsiveness to concurrent radiation and immune checkpoint blockade. Nat Cancer 3, 1498–1512. 10.1038/s43018-022-00467-x

Spurr, L.F., Weichselbaum, R.R., Pitroda, S.P., 2022b. Tumor aneuploidy predicts survival following immunotherapy across multiple cancers. Nat Genet 54, 1782–1785. 10.1038/s41588-022-01235-4

Suter, E.C., Schmid, E.M., Harris, A.R., Voets, E., Francica, B., Fletcher, D.A., 2021. Antibody:CD47 ratio regulates macrophage phagocytosis through competitive receptor phosphorylation. Cell Reports 36, 109587. 10.1016/j.celrep.2021.109587

Taylor, A.M., Shih, J., Ha, G., Gao, G.F., Zhang, X., Berger, A.C., Schumacher, S.E., Wang, C., Hu, H., Liu, Jianfang, The Cancer Genome Atlas Reseach Network, Lazar, A.J., Cherniack, A.D., Beroukhim, R., Meyerson, M., 2018. Genomic and Functional Approaches to Understanding Cancer Aneuploidy. Cancer Cell 33, 676-689.e3. 10.1016/j.ccell.2018.03.007

Tripathi, R., Modur, V., Senovilla, L., Kroemer, G., Komurov, K., 2019. Suppression of tumor antigen presentation during aneuploid tumor evolution contributes to immune evasion. OncoImmunology 8, 1657374. 10.1080/2162402X.2019.1657374

Tsai, R.K., Discher, D.E., 2008. Inhibition of “self” engulfment through deactivation of myosin-II at the phagocytic synapse between human cells. Journal of Cell Biology 180, 989–1003. 10.1083/jcb.200708043

Vasudevan, A., Baruah, P.S., Smith, J.C., Wang, Z., Sayles, N.M., Andrews, P., Kendall, J., Leu, J., Chunduri, N.K., Levy, D., Wigler, M., Storchová, Z., Sheltzer, J.M., 2020. Single-Chromosomal Gains Can Function as Metastasis Suppressors and Promoters in Colon Cancer. Developmental Cell 52, 413–428.e6. 10.1016/j.devcel.2020.01.034

Vasudevan, A., Schukken, K.M., Sausville, E.L., Girish, V., Adebambo, O.A., Sheltzer, J.M., 2021. Aneuploidy as a promoter and suppressor of malignant growth. Nat Rev Cancer 21, 89–103. 10.1038/s41568-020-00321-1

Wang, R.W., Viganò, S., Ben-David, U., Amon, A., Santaguida, S., 2021. Aneuploid senescent cells activate NF-κB to promote their immune clearance by NK cells. EMBO Reports 22. 10.15252/embr.202052032

Willingham, S.B., Volkmer, J.-P., Gentles, A.J., Sahoo, D., Dalerba, P., Mitra, S.S., Wang, J., Contreras-Trujillo, H., Martin, R., Cohen, J.D., Lovelace, P., Scheeren, F.A., Chao, M.P., Weiskopf, K., Tang, C., Volkmer, A.K., Naik, T.J., Storm, T.A., Mosley, A.R., Edris, B., Schmid, S.M., Sun, C.K., Chua, M.-S., Murillo, O., Rajendran, P., Cha, A.C., Chin, R.K., Kim, D., Adorno, M., Raveh, T., Tseng, D., Jaiswal, S., Enger, P.Ø., Steinberg, G.K., Li, G., So, S.K., Majeti, R., Harsh, G.R., van de Rijn, M., Teng, N.N.H., Sunwoo, J.B., Alizadeh, A.A., Clarke, M.F., Weissman, I.L., 2012. The CD47-signal regulatory protein alpha (SIRPa) interaction is a therapeutic target for human solid tumors. Proc. Natl. Acad. Sci. U.S.A. 109, 6662–6667. 10.1073/pnas.1121623109

Xian, S., Dosset, M., Almanza, G., Searles, S., Sahani, P., Waller, T.C., Jepsen, K., Carter, H., Zanetti, M., 2021. The unfolded protein response links tumor aneuploidy to local immune dysregulation. EMBO Rep 22. 10.15252/embr.202152509

Zhou, Y., Fei, M., Zhang, G., Liang, W.-C., Lin, W., Wu, Y., Piskol, R., Ridgway, J., McNamara, E., Huang, H., Zhang, J., Oh, J., Patel, J.M., Jakubiak, D., Lau, J., Blackwood, B., Bravo, D.D., Shi, Y., Wang, J., Hu, H.-M., Lee, W.P., Jesudason, R., Sangaraju, D., Modrusan, Z., Anderson, K.R., Warming, S., Roose-Girma, M., Yan, M., 2020. Blockade of the Phagocytic Receptor MerTK on Tumor-Associated Macrophages Enhances P2X7R-Dependent STING Activation by Tumor-Derived cGAMP. Immunity 52, 357–373.e9. 10.1016/j.immuni.2020.01.014

